# Selective GRAB sensors reveal distinct endocannabinoid dynamics *in vivo*

**DOI:** 10.64898/2026.08.06.743280

**Authors:** Ruyi Cai, Yueqi Yang, Shangxuan Cai, Ana Silva de Sousa, Kathryn L. Todd, Lei Wang, Weijian Teo, Ao Dong, Shuaiyu Chen, Hui Dong, Huan Wang, Zhaofa Wu, Yanling Qiao, Peng Xu, Chen Song, Stephanie J. Cragg, Yulong Li

**Affiliations:** State Key Laboratory of Membrane Biology, School of Life Sciences, Peking University, Beijing 100871, China; PKU-IDG/McGovern Institute for Brain Research, Beijing 100871, China; Peking-Tsinghua Center for Life Sciences, New Cornerstone Science Laboratory, Academy for Advanced Interdisciplinary Studies, Peking University, Beijing 100871, China; Department of Physiology, Anatomy and Genetics, University of Oxford, OX1 3PT, UK; Oxford Parkinson’s Disease Centre, University of Oxford, UK; Aligning Science Across Parkinson’s (ASAP) Collaborative Research Network, Chevy Chase, MD, USA; Center for Quantitative Biology, Academy for Advanced Interdisciplinary studies, Peking University, Beĳing 100871, China; Laboratory of Integrative Physiology, Institute of Genetics and Developmental Biology, Chinese Academy of Sciences, Beijing, China; University of Chinese Academy of Sciences, Beijing, China; Key Laboratory of Drug Monitoring and Control, Drug Intelligence and Forensic Center, Ministry of Public Security, Beijing 100193, China; National Biomedical Imaging Center, Peking University, Beijing 100871, China

## Abstract

The endocannabinoid system modulates diverse physiological processes via two endogenous lipid ligands, 2-arachidonoylglycerol (2-AG) and anandamide (AEA); however, their specific spatiotemporal dynamics remain poorly understood owing to the lack of selective probes. Here, we developed GRAB_2-AG2.0_ and GRAB_AEA2.0_, two genetically encoded fluorescent sensors that selectively detect 2-AG and AEA, respectively. Both sensors exhibited high apparent affinity and molecular specificity for their respective ligands, enabling the real-time detection of 2-AG and AEA release evoked by electrical stimulation in cultured neurons and acute brain slices. In freely behaving mice, these sensors revealed ligand- and context-specific eCB dynamics: aversive stimulation preferentially evoked 2-AG, whereas psychoactive drugs produced distinct 2-AG and AEA responses. Notably, Δ9-THC elicited a sustained 2-AG signal in the nucleus accumbens shell, and local deletion of *Dagla* markedly attenuated both this signal and Δ9-THC-induced hypolocomotion. These sensors therefore enable detecting 2-AG and AEA signaling seperately and reveal an endogenous 2-AG component of the behavioral response to Δ9-THC.

## Introduction

The endocannabinoid system regulates a vast array of neurophysiological processes, including memory, emotion, addiction, and sensory perception^1–4^. Its signaling is mediated primarily by two principal lipid endocannabinoids (eCBs), namely 2-arachidonoylglycerol (2-AG) and *N*-arachidonoylethanolamine (AEA, also known as anandamide), which typically act as retrograde messengers on the cannabinoid receptors CB1R and CB2R^5–9^.

A distinctive feature of the endocannabinoid system is that two chemically different ligands—2-AG and AEA—target the same primary receptors. Despite their shared receptor targets, whether 2-AG and AEA perform redundant or differential physiological roles remains an open question. Although these two ligands possess a similar structural backbone, they are synthesized and degraded via fundamentally distinct enzymatic pathways. Moreover, pharmacological and genetic studies targeting these pathways suggest they may have different functions^10–18^. For example, both 2-AG and AEA appear to regulate anxiety-like behaviors, yet their roles in memory are controversial^12,17,19–21^. Adding to this complexity, some clues indirectly suggest that AEA can activate the ionotropic receptor TRPV1 (transient receptor potential vanilloid 1) and may affect 2-AG levels, suggesting possible cross-regulation between these signaling pathways^22–25^. At the synaptic level, a recent report suggests that 2-AG mediates depolarization-induced suppression of excitation by activating presynaptic CB1Rs, whereas AEA may promote synaptic potentiation via CB1Rs on astrocytes, supporting functional differences between these two eCB ligands^26^.

These putative functional differences between 2-AG and AEA highlight the need for tools that can resolve the individual spatiotemporal dynamics of each ligand. However, progress has been hampered by the limitations associated with existing detection methods^27,28^. For example, direct quantification approaches such as tissue extraction and microdialysis coupled with analytical chemistry have been used in attempts to dissect eCB dynamics in various biological models such as drug intake and addiction^28–35^. However, tissue extraction provides only a static measurement and is susceptible to severe post-mortem artifacts, particularly the rapid accumulation of 2-AG^28^. Moreover, while microdialysis can be used to detect eCBs *in vivo*, its low temporal resolution (typically exceeding 10 minutes) fails to capture transient, sub-second neurotransmission events^6–8,28,36,37^. To address the need to detect eCBs in real time, previously developed GRAB_eCB2.0_ (eCB2.0), a genetically encoded fluorescent sensor, enabled the monitoring of global eCB dynamics^38^. eCB2.0 sensor was developed based on the GPCR activation–based (GRAB) strategy, which has been widely used to develop neurochemical sensors^39–51^. The eCB2.0 sensor has been used to monitor eCB signals in diverse physiological and pathophysiological contexts, including emotional processing, memory, epileptic seizures, and neurodegenerative disease^52–58;^ however, because it uses CB1R as its backbone, it cannot distinguish between 2-AG and AEA^38^. This inability to resolve the individual contributions of these keys signaling molecules remains a critical technological barrier to fully dissecting the complexity of the endocannabinoid system.

This limitation is particularly important for understanding how psychoactive cannabinoids interact with endogenous eCB signaling. Δ9-Tetrahydrocannabinol (Δ9-THC), the major psychoactive constituent of cannabis, is a partial agonist at cannabinoid receptors and produces a characteristic set of acute behavioral effects, including hypolocomotion^59–65^. Although these effects are generally attributed to the direct activation of cannabinoid receptors by Δ9-THC^59–61^, it remains unclear whether Δ9-THC also recruits endogenous 2-AG or AEA signaling and whether this endogenous response contributes to its behavioral actions. Ligand-selective sensors with sufficient temporal resolution are therefore needed to determine how Δ9-THC engages endogenous eCB pathways *in vivo*.

To overcome these barriers, we combined computation-assisted structural analysis with high-throughput screening to develop two genetically encoded fluorescent sensors that selectively detect 2-AG and AEA, respectively. We characterized their selectivity, sensitivity and fluorescence responses in cultured cells and primary neurons. We then used these sensors to compare 2-AG and AEA dynamics during neuronal activity, aversive stimulation and psychoactive drug exposure, and to determine whether Δ9-THC recruits endogenous eCB signaling and whether this signaling contributes to its locomotor effects.

## Results

### AlphaFold3-inspired identification of mutation targets for selective sensors

To engineer a pair of sensors capable of selectively detecting 2-AG or AEA, we employed a structure-guided strategy that integrates computational prediction with high-throughput screening, using the eCB2.0 sensor as the starting scaffold (Fig. 1a). We first predicted the structural models of CB1R in complex with 2-AG or AEA using AlphaFold3^66^, a state-of-the-art deep learning tool with high accuracy in modeling biomolecular complexes. To ensure comprehensive sampling of interaction conformations, we generated 500 independent structures for each CB1R-ligand complex by varying the number of diffusion samples (set as 10), random seeds (number of seeds set as 10), and recycle numbers (set as 0, 1, 2, 3, and 10, respectively) in AlphaFold3 (Fig. 1b). We superimposed the experimentally resolved CB1R–AMG315 (an AEA analog) structure (PDB ID: 8GHV) onto the AlphaFold3-predicted CB1R–2-AG and CB1R–AEA structures. Comparison of the predicted and experimentally determined structures shows a close match, supporting the structural plausibility of the predicted binding poses. (Supplementary Fig. 1a). Subsequent analysis focused on the CB1R residues located within 4 Å of the ligand heavy atoms (‘C’, ‘N’, and ‘O’)—a distance threshold defining putative interacting sites—and quantified the interaction frequency per residue across the 500 sampled structures^67^ (Fig. 1b). We reasoned that residues exhibiting the highest interaction frequencies would be promising engineering targets and thus selected the most frequent sites for each ligand. Some high-contact-frequency residues were common to both 2-AG and AEA: F108^N-term^, F2.57, F2.64, H2.65, T3.33, I267^ECL2^ and W5.43 (Fig. 1C; Supplementary Fig. 1b). In addition, F268^ECL2^, S7.39 showed higher contact frequency with 2-AG (Fig. 1c; Supplementary Fig. 1c), whereas S2.60, F2.61, F3.25, F3.36, and F7.35 were observed to interact more frequently with AEA (Fig. 1c, Supplementary Fig. 1d). Visualizing these complexes with the highlighted high-contact-frequency residues revealed that their side chains orient toward the bound ligands, suggesting functional importance in mediating CB1R-ligand interactions (Supplementary Fig. 1b–d).

**Fig. 1:**
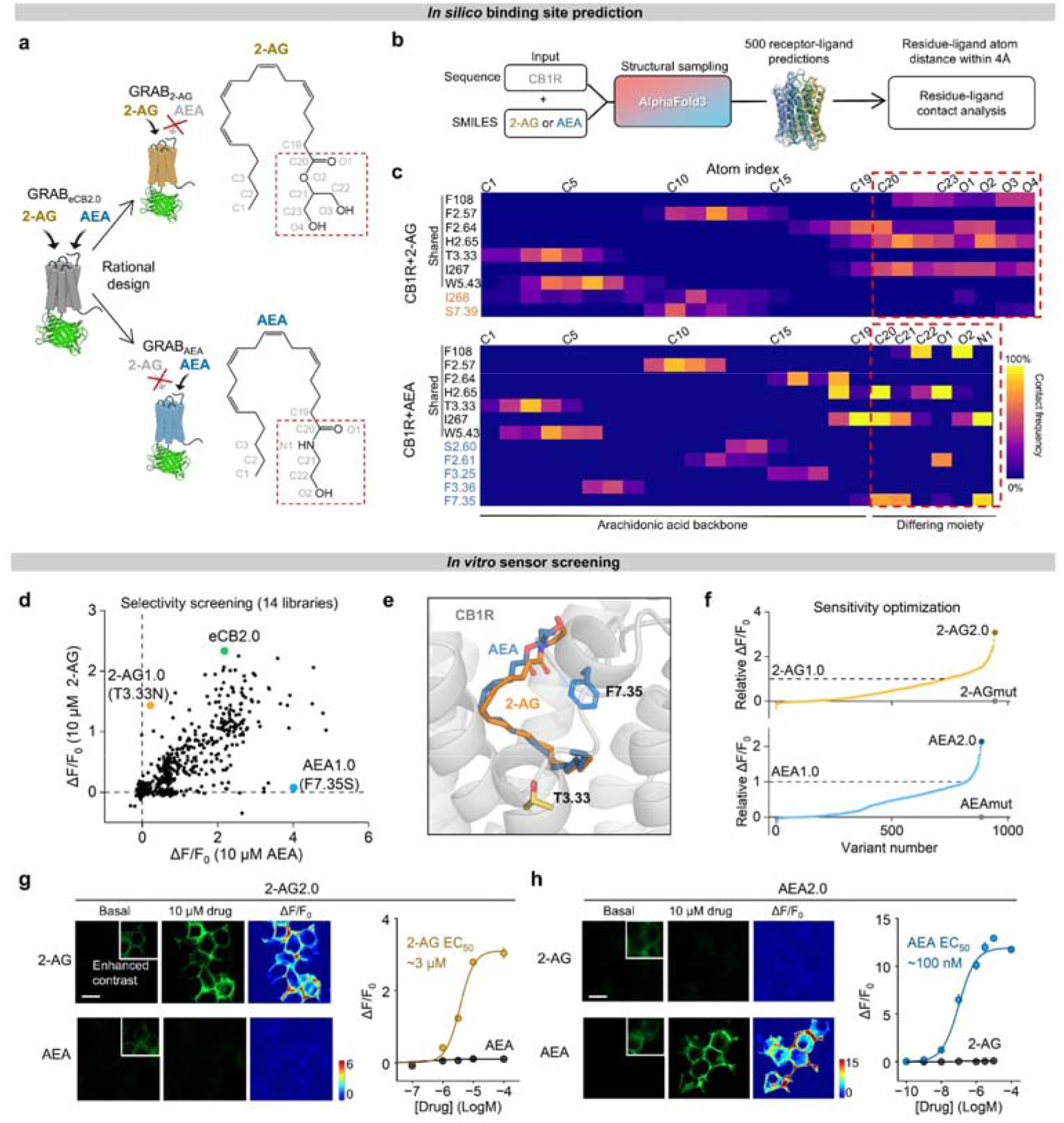
Computation-assisted development of GRAB sensors for selectively detecting 2-AG and AEA. a. Schematic showing the design strategy for engineering 2-AG- and AEA-selective sensors, both derived from the eCB2.0 sensor as the parental backbone. Also shown are the chemical structures of 2-AG and AEA; key structural differences are highlighted in dashed boxes with atom numbering labeled. b. Workflow of the *in silico* calculation pipeline. c. The high-contact-frequency residues of CB1R for 2-AG (top) AEA (bottom) determined based on the AlphaFold3-predicted structures. Residues colored in black, yellow and blue indicate 2-AG and AEA shared, 2-AG unique, and AEA unique high-contact-fequency residues. d, f. Summary of the selectivity screening (d) and sensitivity optimization (f) results. For selectivity screening, each variant’s response (ΔF/F_0_) to 2-AG and AEA is plotted on the *y*- and *x*-axes, respectively. The green circle indicates the eCB2.0 sensor, and the blue and yellow circles indicate the AEA1.0 and 2-AG1.0 sensors, respectively, which were subsequently optimized to produce the AEA2.0 and 2-AG2.0 sensors, respectively (d). For sensitivity optimization, the relative ΔF/F_0_ value is plotted for each variant; the top and bottom panels show optimization of the 2-AG and AEA sensor variants, respectively. The dashed horizontal lines indicate the baseline responses of 2-AG1.0 and AEA1.0 (f). e. Structure of WT CB1R with 2-AG and AEA predicted by AlphaFold3, highlight residues that mutated to develop 2-AG1.0 (T3.33N) and AEA1.0 (F7.35S) sensors. g, h. Left: representative images of sensor expression and fluorescence changes in HEK293T cells expressing 2-AG2.0 (g) or AEA2.0 (h) in response to 100 μM 2-AG or AEA. Scale bars, 20 μm. Right: dose–response curves for 2-AG2.0 (g) and AEA2.0 (h) measured in HEK293T cells, with corresponding EC_50_ values; n = 3 independent experiments. In this and subsequent figures, summary data are presented as the mean ± SEM.

### Screening and optimization of highly selective 2-AG and AEA sensors

Based on our *in silico* predictions, we generated mutation libraries by performing site-saturation mutagenesis at candidate ligand-recognition sites within the eCB2.0 sensor. We then expressed these sensor variants in HEK293T cells and performed high-throughput screening by measuring each variant’s fluorescence change in response to 10 μM 2-AG or 10 μM AEA (Fig. 1d). As expected, the parent eCB2.0 sensor showed limited selectivity and responded robustly to both ligands, with a peak change in fluorescence (ΔF/F_0_) of ∼200%. We then screened approximately 570 variants from 14 mutation libraries and identified two variants with substantially improved specificity compared to eCB2.0 (Fig. 1d). One variant containing the T3.33N mutation had a high response to 2-AG (ΔF/F_0_ ∼150%) with no detectable response to AEA; we named this variant 2-AG1.0 (Fig. 1d, e). Conversely, one variant containing the F7.35S mutation responded robustly to AEA (ΔF/F_0_ ∼400%) with only a negligible response to 2-AG; we named this variant AEA1.0 (Fig. 1d, e).

Next, we optimized these two prototype sensors by screening ∼1000 additional variants for each sensor, targeting GPCR activation–related residues, the linkers connecting the GPCR with cpEGFP, and residues within the cpEGFP moiety. This optimization substantially improved each sensor’s sensitivity. For 2-AG1.0, combining 10 mutations (E5.37R and T7.39S in the GPCR; M308N, R311H, T312P, and R570K in the linker regions; and I23G, S96Q, T140S, and N202T in cpEGFP) yielded 2-AG2.0, which exhibited a 3-fold higher increase in ΔF/F_0_ compared to 2-AG1.0; introducing F2.64A produced a non-responsive 2-AG control sensor, 2-AGmut (Fig. 1f top; Supplementary Fig. 2a, b). Similarly, introducing 11 mutations (E5.37K, T7.39L, and W6.48F in the GPCR; M308N, R311H, T312P, and R570K in the linker regions; and I23G, S96Q, T140S, and N202Y in cpEGFP) in AEA1.0 resulted in AEA2.0, which exhibited an approximately 2-fold higher increase in ΔF/F_0_ compared to AEA1.0; introducing F2.64A produced the AEA control sensor, AEAmut (Fig. 1f bottom; Supplementary Fig. 2a, c).

### *In vitro* characterization of the 2-AG2.0 and AEA2.0 sensors

We then characterized 2-AG2.0 and AEA2.0 in HEK293T cells and cultured rat primary cortical neurons. When expressed in HEK293T cells, both sensors localized primarily to the plasma membrane and exhibited a robust ligand-evoked increase in fluorescence with high selectivity and appropriate apparent affinity for their respective ligand (Fig. 1g, h). Specifically, 2-AG2.0 responded to 2-AG in a dose-dependent manner with an EC_50_ of 3 μM and a peak ΔF/F_0_ of ∼300% in response to 100 μM 2-AG treatment; in contrast, AEA elicited no significant response in 2-AG2.0-expressing cells at any concentration tested (Fig. 1g). Similarly, AEA2.0 responded to AEA in a dose-dependent manner, with an EC_50_ of 100 nM (similar to the wild-type CB1R; data not shown) and a ∼1200% peak ΔF/F_0_; in contrast, 2-AG elicited no significant response in AEA2.0-expressing cells at any concentration tested (Fig. 1h).

We next characterized these sensors’ properties in cultured primary rat cortical neurons by adeno-associated virus (AAV) infection. Consistent with our results obtained with HEK293T cells, both sensors showed good membrane expression (Fig. 2a, b), robust responses (Fig. 2a, b, c, f), similar apparent affinity values (Fig. 2c, f), and high selectivity for their respective ligands (Fig. 2c, f). Specifically, 2-AG2.0 had a peak ΔF/F_0_ of ∼600% in response to 100 μM 2-AG, an EC_50_ of 1.4 μM, and no significant response to AEA within its reported physiological range (picomolar to nanomolar)^28^ (Fig. 2c). Moreover, AEA2.0 had a peak ΔF/F_0_ of ∼1600% in response to 100 μM AEA, an EC_50_ of 90 nM, and no significant response to 2-AG at concentrations relevant to physiological brain levels^28^ (Fig. 2f).

**Fig. 2:**
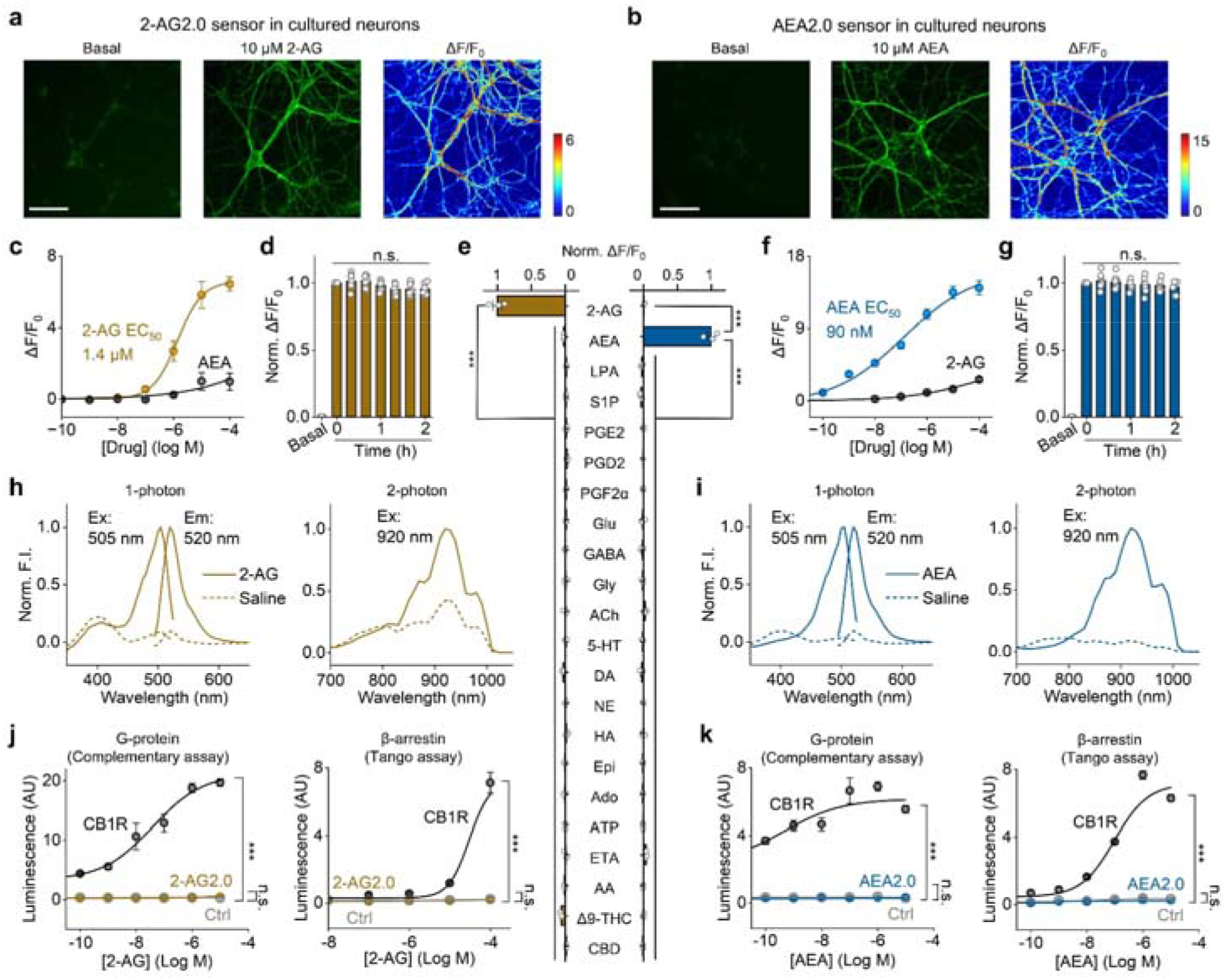
*In vitro* characterization of 2-AG2.0 and AEA2.0 a, b. Representative fluorescence images and fluorescence change in response to 2-AG and AEA in cultured cortical neurons expressing 2-AG2.0 (a) or AEA2.0 (b). Scale bar, 100 μm. c, f. Dose–response curves measured in neurons expressing 2-AG2.0 (c) or AEA2.0 (f), with corresponding EC_50_ values; n = 4 cultures each. d, g. Summary of the change in 2-AG2.0 (d) and AEA2.0 (g) fluorescence normalized to peak fluorescent response in neurons exposed continuously for 2 h to MDMB-4en-PINACA (1 μM); n = 7 cultures each. e. Normalized fluorescence changes in response to the indicated compounds (each applied at 10 μM) measured in cells expressing 2-AG2.0 (indicated in yellow) or AEA2.0 (indicated in blue); n = 3 cultures/group. LPA, lysophosphatidic acid; S1P, sphingosine-1-phosphate; PGE2, prostaglandin E2; PGD2, prostaglandin D2; PGF2α, prostaglandin F2α; Glu, glutamate; GABA, γ-Aminobutyric acid; Gly, glycine; ACh, acetylcholine; 5-HT, 5-hydroxytryptamine; DA, dopamine; NE, norepinephrine; HA, histamine; Epi, epinephrine; Ado, adenosine; ATP, adenosine triphosphate; ETA, ethanolamine; AA, arachidonic acid; Δ9-THC, Δ9-tetrahydrocannabinol; CBD, cannabidiol. n = 3 cultures for each group. h, i. One-photon excitation (Ex) and emission (Em) spectra (left) and 2-photon excitation spectra (right) measured in HEK293T cells expressing 2-AG2.0 (h) or AEA2.0 (i) in saline and in the presence of 2-AG or AEA. The excitation and emission peaks are labeled. F.I., fluorescence intensity. j, k. The luciferase complementation assay (left) and Tango assay (right) were used to assess downstream coupling to G_i_ proteins and β-arrestin, respectively, in HEK293T or HTLA cells expressing WT CB1R or either 2-AG2.0 (j) or AEA2.0 (k). Ctrl refers to cells expressing LgBit alone (for the luciferase complementation assay) or no receptor (for the Tango assay); n = 3 cultures each.

We then tested the specificity of these two sensors using a panel of bioactive lipids, neurotransmitters, neuromodulators, eCB degradation products, and major phytocannabinoids when expressed in HEK293T cells. Neither sensor had a detectable response to any of the tested compounds other than its respective ligand, indicating high molecular specificity and low potential for off-target activation (Fig. 2e). Notably, both sensors had minimal response to Δ9-THC, suggesting that the sensors can be used in phytocannabinoid-related studies (Fig. 2e; Supplementary Fig. 3a, c). We also tested seven synthetic cannabinoids and found differential response profiles between 2-AG2.0 and AEA2.0, although 4CN-CUMYL-BUTINACA, 5F-ADB, and MDMB-4en-PINACA activated both sensors (Supplementary Fig. 3b, d). Furthermore, when expressed in cultured neurons, both sensors were highly stable, with no significant change in peak fluorescence during 2 hours of continuous exposure to 1 μM MDMB-4en-PINACA, suggesting that the sensors do not undergo β-arrestin-mediated internalization (Fig. 2d, g). Moreover, the sensors had similar spectral properties, with 1-photon excitation and emission peaks at ∼505 nm and ∼520 nm, respectively, and 2-photon excitation peaks at ∼920 nm (Fig. 2h, i). Finally, we assessed the sensors’ potential for downstream coupling using the mini-G protein complementary bioluminescent assay (for G_i_) and the Tango assay (for β-arrestin). Compared to the wild-type (WT) CB1R, both sensors showed no detectable coupling to either the G_i_ or β-arrestin pathways, indicating that their expression has minimal effect on endogenous intracellular signaling (Fig. 2j, k).

### Activity-dependent release of endogenous 2-AG in cultured neurons measured using 2-AG2.0

Next, we tested whether these sensors can be used to report eCB release in response to depolarization in cultured cortical neurons. Using the non-selective eCB2.0 sensor combined with pharmacological validation, we previously reported that 2-AG, rather than AEA, is likely the primary eCB released from cortical neurons upon electrical stimulation^38^. To test this directly, we loaded cultured neurons expressing either 2-AG2.0 or AEA2.0 with the red fluorescent calcium indicator Calbryte 590-AM (Fig. 3a). Electrical stimulation applied at 20 Hz elicited robust calcium transients in both 2-AG2.0-and AEA2.0-expressing neurons, but only the cells expressing 2-AG2.0 had an increase in green fluorescence (Fig. 3b–g). Consistent with our previous findings^38^, a 100-pulse train evoked a large response in 2-AG2.0-expressing neurons (ΔF/F_0_ ∼100%), with slower kinetics than the calcium signal (Fig. 3b, d); in contrast, cells expressing AEA2.0 had no detectable response, despite a robust calcium signal (Fig. 3c, f). To confirm sensor expression, we perfused the neurons with 10 µM 2-AG or AEA and measured a robust fluorescence increase in the corresponding sensors (Fig. 3e, g). We then tested a range of stimulus intensities ranging from 5 to 100 pulses applied at 20 Hz and found pulse number-dependent increases in both the calcium and 2-AG2.0 signals in neurons expressing 2-AG2.0 (Fig. 3d, right), but only an increase in the calcium signal in cells expressing AEA2.0 (Fig. 3f, right). Quantification of peak ΔF/F_0_ in cells expressing 2-AG2.0 revealed a strong direct correlation between the 2-AG2.0 and calcium signals (Pearson’s *r*=0.97, *P*=0.0075), but no correlation in cells expressing AEA2.0 (Pearson’s *r*=-0.70, *P*=0.19).

**Fig. 3:**
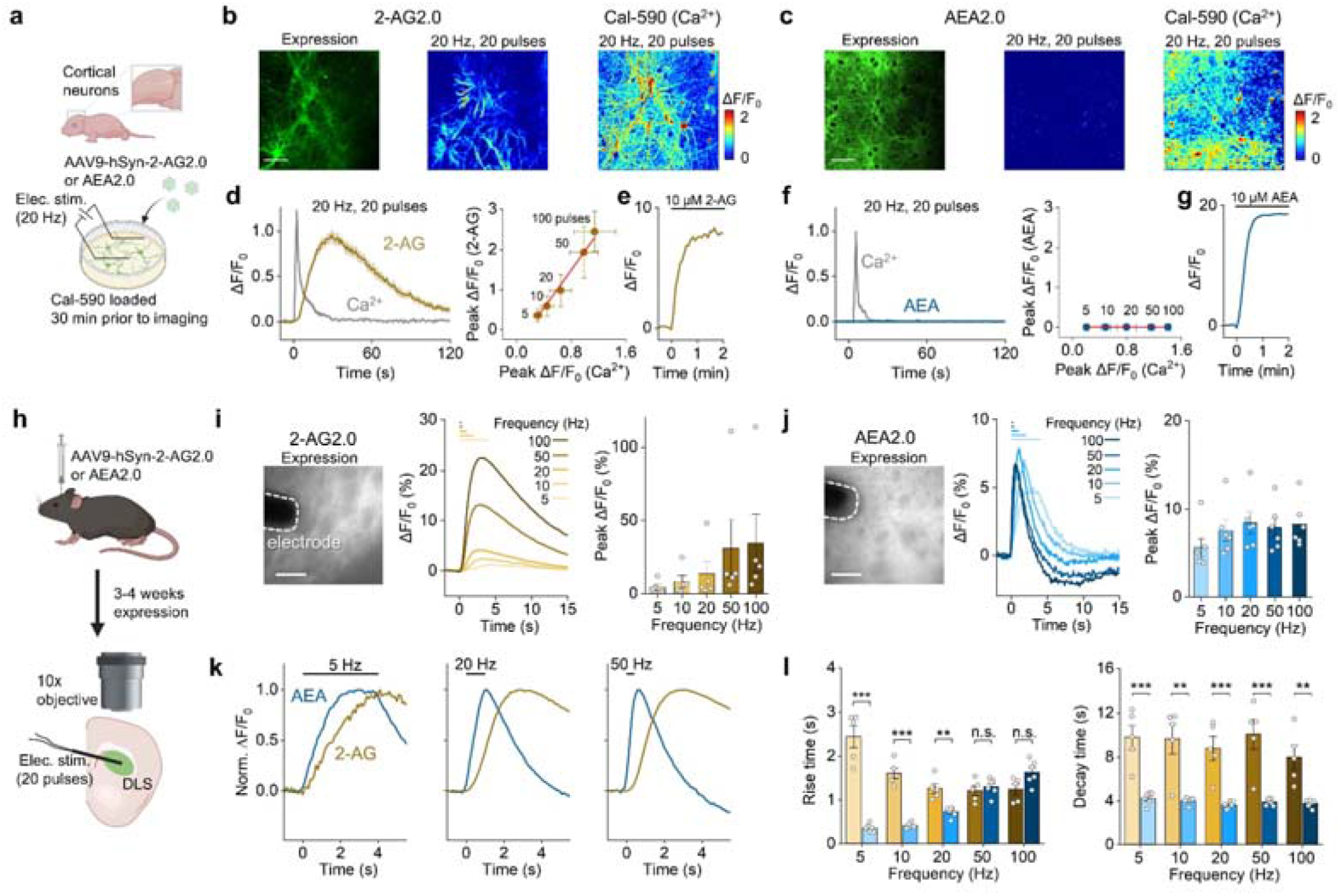
The 2-AG2.0 and AEA2.0 sensors can report the activity-dependent release of 2-AG and AEA, respectively, both *in vitro* and *ex vivo* a. Schematic diagram depicting the experimental design for electrical stimulation of cultured rat cortical neurons. b, c. Representative fluorescence images (left) and fluorescence responses (middle) in cortical neurons expressing 2-AG2.0 (b) or AEA2.0 (c) and loaded with the calcium indicator dye Calbryte 590-AM, stimulated with 20 electrical pulses applied at 20 Hz (right). Scale bar, 100 μm. d, f. Fluorescence traces of 2-AG2.0 (d) or AEA2.0 (f) and calcium signals during electrical stimulation applied at time 0 (left), and the correlation between the peak calcium response and peak 2-AG2.0 (d) or AEA2.0 (f) response following the indicated number of pulses (right). Pearson’s correlation for 2-AG2.0 and calcium: *r* =0.97, *P* =0.0075. Pearson’s correlation for AEA2.0 and calcium: *r* = −0.70, *P* = 0.19. e, g. Representative fluorescence traces of 2-AG2.0 (e) and AEA2.0 (g); the horizontal bars indicate the bath perfusion of 2-AG or AEA. h. Schematic diagram depicting the experimental design for acute brain slice imaging and electrical stimulation. i, j. Left: representative fluorescence images showing expression of 2-AG2.0 (i) or AEA2.0 (j) in the dorsolateral striatum (DLS). Middle: representative traces of 2-AG2.0 (i) and AEA2.0 (j) fluorescence; at time 0, 20 electrical pulses were applied at 5, 10, 20, 50, or 100 Hz. Right: summary of the peak fluorescence changes in 2-AG2.0 (i) and AEA2.0 (j); n = 5–6 slices per group. k. Representative traces of 2-AG2.0 (yellow) and AEA2.0 (blue) fluorescence responses evoked by 20 pulses applied at 5 Hz (left), 10 Hz (middle), and 20 Hz (right). l. Rise and decay times (10–90%) of 2-AG2.0 (yellow bars) and AEA2.0 (blue bars) fluorescence plotted as a function of stimulation frequency (20 pulses applied at 5, 10, 20, 50, or 100 Hz); n = 5–6 slices per group.

### Detecting endogenous 2-AG and AEA release in acute brain slices

We next expressed the sensors in a more physiologically relevant preparation, acute brain slices, in order to monitor 2-AG and AEA dynamics *ex vivo*. We injected AAVs expressing either 2-AG2.0 or AEA2.0 under the human synapsin (hSyn) promoter into the dorsolateral striatum (DLS), a CB1R-rich region^68,69^, and prepared acute brain slices 3–4 weeks later as described previously^70,71^ (Fig. 3h). We first applied increasing pulse numbers (5–100 pulses) at 20 Hz and found that both 2-AG2.0 and AEA2.0 responded in a pulse number-dependent manner (with peak ΔF/F_0_ of ∼5–20% and ∼5–15%, respectively) (Supplementary Fig. 4a, b, d, e). The rise and decay time of both sensors increased in response to increasing pulse numbers (Supplementary Fig. 4b, e). Interestingly, when we applied 20 pulses at increasing frequencies (5–100 Hz), we found differences between 2-AG2.0 and AEA2.0. Specifically, the 2-AG2.0 signal was highly frequency-dependent, with peak ΔF/F_0_ increasing from ∼2% to 20% as frequency increased (Fig. 3i); in contrast, the peak AEA2.0 response was ∼7% regardless of frequency (Fig. 3j). We also found differences in kinetics between the 2-AG2.0 and AEA2.0 responses, with the AEA2.0 signal having significantly faster rise and decay times compared to the 2-AG2.0 signal, suggesting differences in kinetics between these two chemicals (Fig. 3k, l; Supplementary Fig. 4g). Bath-application of exogenous ligands confirmed sensor expression and selectivity in these slices (Supplementary Fig. 4c, f).

### The 2-AG2.0 sensor reveals 2-AG release in the basolateral amygdala and the nucleus accumbens in response to foot shock in freely moving mice

Having validated the sensors in acute slices, we investigated whether 2-AG and/or AEA are released *in vivo* during specific behavioral paradigms. We focused on two CB1R-rich regions involved in valence processing, namely the basolateral amygdala (BLA) and the nucleus accumbens (NAc) core^69,72^. We previously showed that foot shock induces robust endocannabinoid release in the BLA using the non-selective eCB2.0 sensor^38^ (Fig. 4a, b); however, the specific lipid(s) involved in this response is currently unknown. To address this question, we expressed 2-AG2.0 and AEA2.0 in the BLA in opposing hemispheres and recorded the change in fluorescence using fiber photometry in freely moving WT mice (Fig. 4c). We found that foot shock evoked a rapid, transient increase in 2-AG2.0 fluorescence (Fig. 4e, left) that was reminiscent of the response measured for eCB2.0 (Fig. 4b). In contrast, foot shock failed to induce a response in AEA2.0 in the BLA (Fig. 4e, middle); as a control to confirm AEA2.0 expression, an i.p. injection of MDMB-4en-PINACA elicited a response in both hemispheres (Supplementary Fig. 5).

**Fig. 4:**
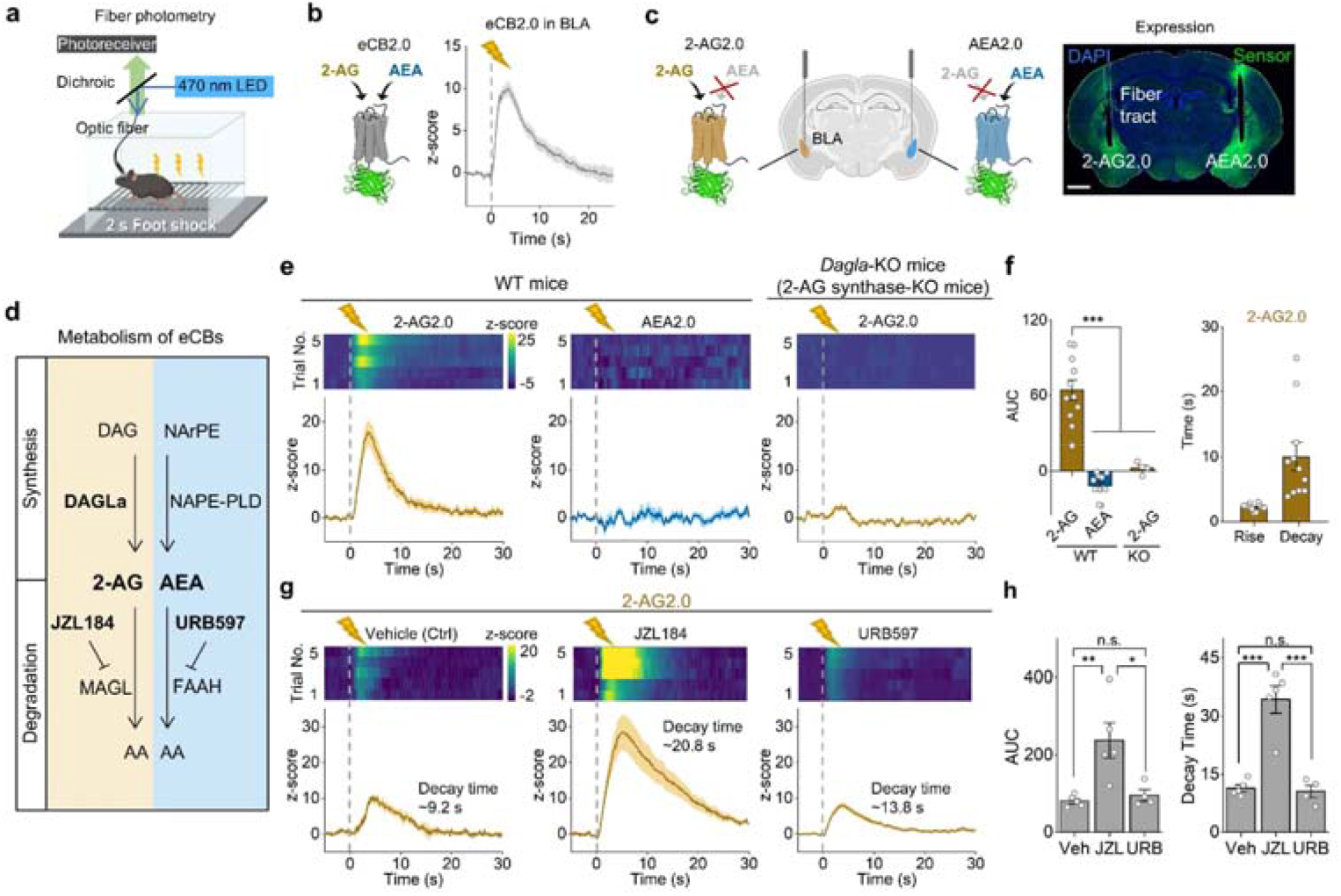
The 2-AG2.0 and AEA2.0 sensors reveal distinct patterns of 2-AG and AEA signaling in response to foot shock. a. Schematic diagram depicting the experimental design for *in vivo* fiber photometry recordings during foot shock in mice expressing either eCB2.0 or 2-AG2.0 and AEA2.0 in opposing hemispheres. b. Left: schematic diagram of the eCB2.0 sensor. Right: example trace of eCB2.0 fluorescence in response to foot shock applied at time 0 (replotted from ^38^). c. Left: schematic diagram depicting the expression of 2-AG2.0 and AEA2.0 in the BLA in opposing hemispheres of the same mouse. Right: histological verification of sensor expression and fiber placement in the BLA; the nuclei were counterstained with DAPI. Scale bar, 1 mm. d. Schematic diagram showing the biosynthetic (top) and degradative (bottom) pathways of AEA and 2-AG; showing the inhibitors for AEA and 2-AG degradation. DAG, diacylglycerol; NArPE, N-arachidonoyl phosphatidylethanolamines; DAGLa, diacylglycerol lipase alpha; NAPE-PLD, N-acyl phosphatidylethanolamine-hydrolyzing phospholipase D; MAGL, monoacylglycerol lipase; FAAH, fatty acid amide hydrolase; AA, arachidonic acid; JZL184, MAGL inhibitor; URB597, FAAH inhibitor. e. Representative pseudocolor heatmaps showing sensor responses over 5 trials (top) and corresponding average fluorescence traces (bottom) measured in the BLA of a WT mouse (left and middle) and *Dagla*-KO (e) mouse expressing 2-AG2.0 (yellow traces) and AEA2.0 (blue traces) in opposing hemispheres. Where indicated, a 2 s foot shock was applied. f. Group summary of the area under the curve (AUC, 0–30 s after shock) (left) for the 2-AG2.0 and AEA2.0 fluorescence signals, and rise and decay time (10–90%) (right) for the 2-AG2.0 fluorescence signals measured in the BLA of WT and Dagla-KO mice following foot shock; n=4–11 mice per group. g. Representative pseudocolor heatmaps showing the responses of 2-AG2.0 over 5 trials (top) and corresponding average fluorescence traces (bottom) measured in the BLA of WT mice expressing 2-AG2.0 following intraperitoneal an (i.p.) injection of vehicle (left), JZL184 (middle), or URB597 (right). Where indicated, a 2-s foot shock was applied. The decay times are indicated in plots. h. Summary of AUC (0–30 s after shock) (left) and decay time (10–90%) (right) for the 2-AG2.0 fluorescence signals measured in the BLA of WT mice treated with an i.p. injection of vehicle (Veh), JZL184 (JZL), or URB597 (URB); n = 4–5 mice per group.

We then confirmed that the signal measured in the BLA reflects 2-AG release using both genetic and pharmacological manipulations. When we expressed 2-AG2.0 and AEA2.0 in mice lacking diacylglycerol lipase alpha (DAGLa, the principal enzyme involved in synthesizing 2-AG), foot shock failed to evoke a signal (Fig. 4d–f). Moreover, in WT mice expressing 2-AG2.0 and AEA2.0 in the BLA, inhibiting MAGL (monoacylglycerol lipase, the enzyme responsible for degrading 2-AG) with JZL184 (16 mg/kg) significantly increased the peak 2-AG2.0 signal and slowed the signal decay induced by foot shock, whereas inhibiting FAAH (fatty acid amide hydrolase, the enzyme responsible for breaking down AEA) with URB597 (0.3 mg/kg) had no effect on the 2-AG2.0 signal (Fig. 4d, g, h). Similar results were obtained when we expressed 2-AG2.0 and AEA2.0 in the NAc core (Supplementary Fig. 6a–d). Interestingly, when we compared the foot shock-evoked 2-AG signals between the BLA and NAc core, we found similar rising kinetics, while the signal measured in the BLA was significantly larger and decayed more rapidly (Supplementary Fig. 6e, f), suggesting brain region-specific differences in 2-AG signaling during an aversive stimulus.

### Nicotine selectively evokes AEA in the NAc shell

The endocannabinoid system is a key modulatory target for addictive drugs, yet few studies have resolved the real-time dynamics of specific endocannabinoids during drug exposure. To examine these dynamics, we expressed 2-AG2.0 and AEA2.0 in the NAc shell, a key hub in addiction-related circuitry^2^, and monitored their fluorescence using fiber photometry (Fig. 5a,b). We first examined the effects of nicotine, cocaine and ethanol. Control experiments in HEK293T cells confirmed that neither nicotine nor cocaine directly activated either sensor (Fig. 5c).

**Fig. 5:**
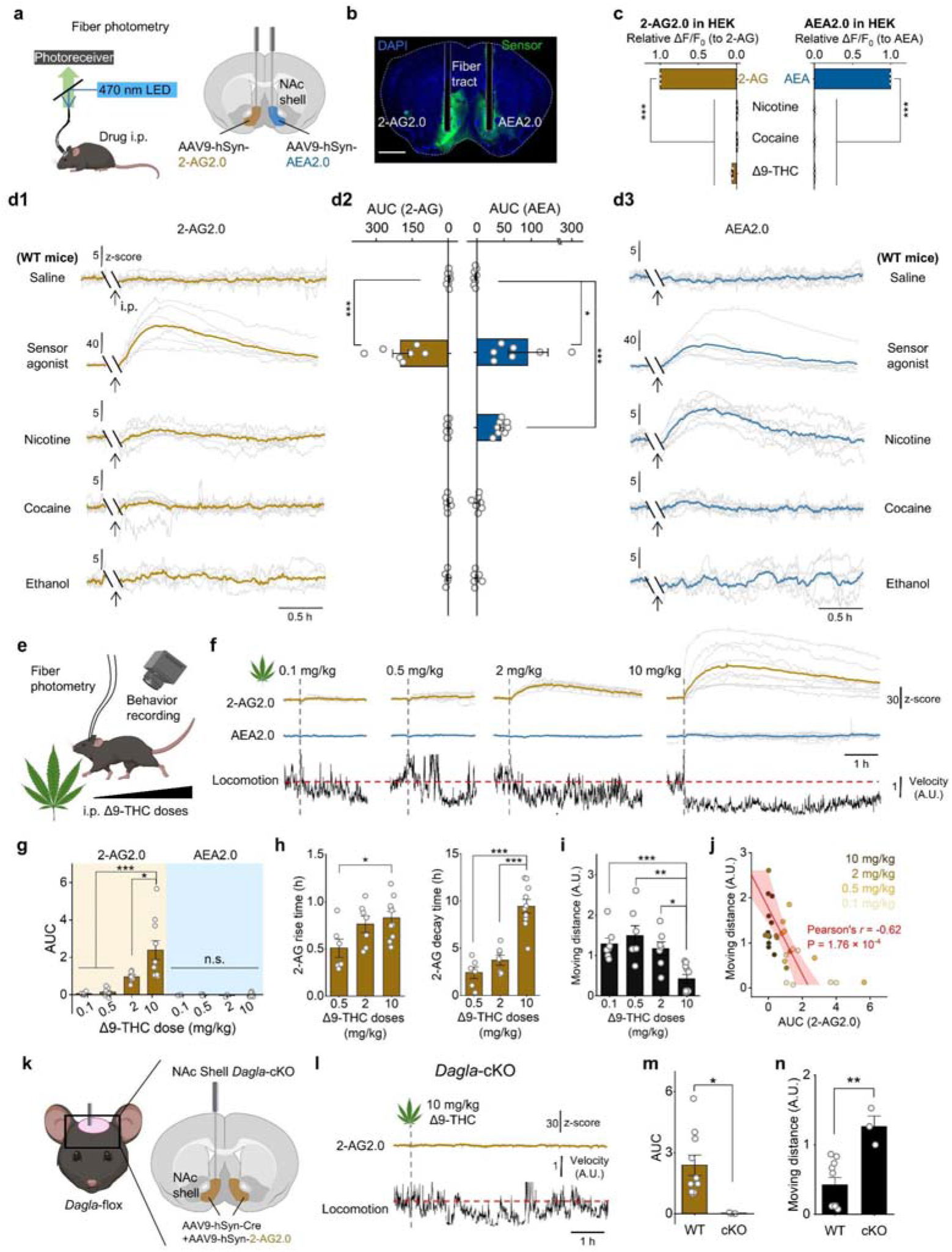
Psychoactive substances differentially mobilize 2-AG and AEA in the nucleus accumbens shell. a. Schematic diagram depicting the experimental strategy for *in vivo* fiber photometry recordings in the mouse NAc shell (for panels B and D1–D3). b. Histological verification of 2-AG2.0 and AEA2.0 expression and fiber placement in the NAc shell. The nuclei were counterstained with DAPI. Scale bar, 1 mm. c. Summary of the 2-AG2.0 (left) and AEA2.0 (right) response to nicotine, cocaine, and Δ9-THC (10 μM) in HEK293T cells (all data are expressed relative to the response the sensor’s respective ligand); n=3 cultures per group. d1, d3. Average fluorescence traces (*z*-scores) of 2-AG2.0 (yellow; d1) and AEA2.0 (blue; d3) recorded in the NAc shell of WT mice before and after i.p. injection of saline, the sensor agonist MDMB-4en-PINACA (0.1 mg/kg), nicotine (0.5 mg/kg), cocaine (20 mg/kg) or ethanol (2 g/kg). The arrows indicate the time of injection; gray lines represent individual mice and colored lines represent the group means. d2, Summary of the area under the curve (AUC; 5–35 min after injection) for the 2-AG2.0 (left) and AEA2.0 (right) signals shown in d1 and d3; n = 5–8 mice per group. e, Schematic depicting simultaneous fiber photometry and open-field locomotor recordings following administration of increasing doses of Δ9-THC. f, 2-AG2.0 (yellow) and AEA2.0 (blue) fluorescence traces and average locomotor-velocity traces following i.p. injection of Δ9-THC at 0.1, 0.5, 2 or 10 mg/kg. Vertical dashed lines indicate the time of injection. Gray lines represent individual fluorescence traces and colored lines represent the group means. n = 10 mice. g. Summary of AUC (5–125 min after injection) of the fluorescence signals measured for 2-AG2.0 and AEA2.0 in response to an i.p. injection of the indicated doses of Δ9-THC; n = 3–10 mice per group. h. Summary of the rise time (left) and decay time (10–90%, right) of the 2-AG2.0 signals measured in response to the indicated doses of Δ9-THC; n = 6–10 mice per group. i. Moving distance from 5 to 125 min after administration of the indicated doses of Δ9-THC. For each mouse, speed was normalized to its mean speed during the pre-injection baseline (−65 to −5 min), and moving distance was calculated by integrating the resulting relative-speed trace over the indicated post-injection window. The resulting values are therefore expressed in arbitrary units (A.U.); n = 7–10 mice per group. j. Moving distance from 5 to 125 min after Δ9-THC injection plotted against the corresponding 2-AG2.0 AUC from 5 to 125 min after injection. Colors indicate the administered Δ9-THC doses. The Pearson correlation coefficient and P value are shown. k. Schematic depicting bilateral co-expression of Cre recombinase and 2-AG2.0 in the NAc shell of *Dagla*-flox mice, generating NAc-shell *Dagla*-cKO mice. l. 2-AG2.0 fluorescence trace (top) and the average locomotor-velocity trace (bottom) recorded in *Dagla*-cKO mice following i.p. injection of Δ9-THC (10 mg/kg). The vertical dashed line indicates the time of injection; gray lines represent individual fluorescence traces and the yellow line represents the group mean. m. AUC of the 2-AG2.0 signal from 5 to 125 min after Δ9-THC injection in WT (replotted from g) and *Dagla*-cKO mice. n = 3–10 mice per group. n. Moving distance from 5 to 125 min after Δ9-THC injection in WT (replotted from i) and *Dagla*-cKO mice, calculated by integrating the relative-speed trace normalized to the pre-injection baseline as described in i. n = 3–10 mice per group.

In WT mice, saline injection elicited no detectable change in either sensor, whereas injection of the sensor agonist MDMB-4en-PINACA (0.1 mg/kg), which activates both sensors, produced robust increases in 2-AG2.0 and AEA2.0 fluorescence (Fig. 5d1–d3). Among the drugs examined, nicotine (0.5 mg/kg), the primary psychoactive component of tobacco, selectively increased AEA2.0 fluorescence without producing a detectable change in the 2-AG2.0 signal (Fig. 5d1–d3). This nicotine-evoked AEA2.0 signal was nearly absent in *Napepld*-KO mice, which lack N-acyl phosphatidylethanolamine-hydrolyzing phospholipase D, an enzyme involved in AEA synthesis (Supplementary Fig. 7a–c), confirming that the nicotine-evoked AEA2.0 signal reflects NAPE-PLD-dependent AEA mobilization in the NAc shell. In contrast, neither cocaine (20 mg/kg) nor ethanol (2 g/kg) produced a detectable change in either 2-AG2.0 or AEA2.0 fluorescence (Fig. 5d1–d3).

### NAc-shell 2-AG synthesis contributes to **Δ**9-THC-induced hypolocomotion

We next asked whether Δ9-THC recruits endogenous 2-AG or AEA signaling in the NAc shell. Because both sensors are engineered from CB1R, we first confirmed that Δ9-THC produced minimal direct activation of either sensor in HEK293T cells (Fig. 5c; Supplementary Fig. 3a,c). We then simultaneously monitored 2-AG2.0 or AEA2.0 fluorescence and open-field locomotion following administration of increasing doses of Δ9-THC (0.1, 0.5, 2 and 10 mg/kg; Fig. 5e,f). Consistent with previous reports^59,62–65^, Δ9-THC produced a dose-dependent reduction in locomotor activity (Fig. 5f,i). In parallel, Δ9-THC elicited a dose-dependent increase in the 2-AG2.0 signal, with progressively longer rise and decay times, whereas the AEA2.0 signal remained unchanged across the doses tested (Fig. 5f–h). Across all doses, the 2-AG2.0 signal was inversely correlated with the distance traveled after Δ9-THC injection (Pearson’s r = −0.62, P = 1.76 × 10^−4^; Fig. 5j), establishing a association between 2-AG mobilization and hypolocomotion.

To determine whether locally synthesized 2-AG contributes to the Δ9-THC-evoked signal and hypolocomotion, we bilaterally co-expressed Cre recombinase and 2-AG2.0 in the NAc shell of *Dagla*-flox mice, thereby generating NAc-shell *Dagla*-cKO mice (Fig. 5k). In these mice, Δ9-THC (10 mg/kg) produced only a minimal increase in 2-AG2.0 fluorescence compared with WT mice (Fig. 5l,m). The reduction in locomotor activity following Δ9-THC injection was also markedly attenuated in *Dagla*-cKO mice (Fig. 5l,n). As a positive control, MDMB-4en-PINACA still elicited a robust 2-AG2.0 response in *Dagla*-cKO mice, confirming sensor expression and responsiveness in the NAc shell (Supplementary Fig. 8a,b). Together, these results show that DAGLα-mediated 2-AG synthesis in the NAc shell is required for the hypolocomotor effect of Δ9-THC.

## Discussion

Our understanding of the endocannabinoid system has long been hindered by a lack of tools capable of resolving the dynamics of the two primary lipid messengers in real time. Here, we addressed this knowledge gap by developing GRAB_2-AG2.0_ and GRAB_AEA2.0_, two genetically encoded sensors that selectively report the release of 2-AG and AEA, respectively. Using these sensors, we provide direct *in vitro*, *ex vivo*, and *in vivo* evidence that 2-AG and AEA have distinct release patterns and dynamics in various biological contexts, indicating that these two eCB ligands may be differentially regulated in a diverse range of physiological and pathological processes.

Our *ex vivo* results reveal fundamental differences in kinetics between cultured neurons and brain slices. The apparent difference between *in vitro* and *ex vivo* preparations may reflect differences in network connectivity, local circuit organization, and/or baseline activity.

Furthermore, within brain slices, we observed distinct release dynamics for 2-AG and AEA. The magnitude of 2-AG release was highly frequency-dependent, increasing with stimulation frequency; in contrast, the magnitude of AEA release was relatively constant regardless of stimulation frequency. Conversely, the AEA signal showed markedly faster rise and decay kinetics compared to 2-AG under the same stimulation conditions; this difference suggests distinct regulatory mechanisms of 2-AG and AEA. Previous studies reported that eCB-mediated short-term and long-term plasticity may be induced by specific neuronal stimulation protocols^6–8,73,74^. Our observation that the magnitude of the 2-AG signal increases with stimulation frequency while the AEA signal remains constant may reflect distinct activity thresholds for mobilizing these distinct ligands, potentially explaining their differential engagement in specific forms of synaptic plasticity.

In freely behaving mice, we resolved the molecular identity of foot shock-evoked eCB dynamics in BLA, showing that 2-AG is the principal eCB involved in these processes. Furthermore, we found that phasic 2-AG release is conserved in the NAc core, suggesting a universal role for 2-AG in buffering acute circuit hyperactivity in response to aversive stimuli.

Our *in vivo* drug-administration experiments further demonstrate that specific psychoactive substances engage distinct eCB pathways. Nicotine selectively mobilized AEA in the NAc shell, consistent with previous reports that nicotine exposure alters AEA levels in reward-related brain regions^29,34^. This finding may also relate to the reported ability of nicotine to facilitate striatal eCB-mediated long-term depression, a form of synaptic plasticity in which AEA has been implicated as a primary mediator^75^. By contrast, cocaine and ethanol did not produce detectable changes in either sensor under the conditions tested, further illustrating that psychoactive drug exposure does not uniformly recruit 2-AG and AEA signaling.

A separate finding was that Δ9-THC elicited a large and sustained 2-AG signal in the NAc shell without producing a detectable AEA signal. More importantly, local deletion of *Dagla* markedly attenuated both the Δ9-THC-evoked 2-AG signal and the accompanying reduction in locomotor activity. These findings indicate that acute Δ9-THC-induced hypolocomotion is not explained solely by the direct action of Δ9-THC at cannabinoid receptors, but depends substantially on DAGLα-mediated 2-AG synthesis in the NAc shell. Because Δ9-THC is generally considered a partial agonist at cannabinoid receptors whereas 2-AG acts as a full agonist^60,61^, one possibility is that Δ9-THC-induced 2-AG release amplifies or prolongs cannabinoid-receptor signaling. This mechanistic interpretation remains a hypothesis, and future experiments combining eCB dynamics imaging with receptor-specific and cell-type-specific manipulations will be needed to determine how Δ9-THC recruits 2-AG synthesis.

In summary, we developed two selective genetically encoded sensors that enable the dynamics of 2-AG and AEA to be resolved independently *in vitro*, *ex vivo* and *in vivo*. These sensors revealed ligand- and context-specific eCB signaling during neuronal stimulation, aversive experience and psychoactive drug exposure. Beyond distinguishing the dynamics of the two endogenous ligands, the sensors uncovered a sustained Δ9-THC-evoked 2-AG signal in the NAc shell and identified local DAGLα-dependent 2-AG synthesis as a major contributor to Δ9-THC-induced hypolocomotion. Thus, this work provides both a new set of tools for eCB detection and evidence that an exogenous cannabinoid can recruit endogenous eCB signaling to shape its behavioral effects.

## Methods

### Animals

All animal experiments were performed in accordance with protocols approved by the Laboratory Animal Care and Use Committees of Peking University and the Animals in Scientific Procedures Act 1986 (amended in 2012) with ethics approval from the University of Oxford, and under the authority of a project license granted by the UK Home Office. Cultured cortical neurons were dissociated from newborn (P0) Sprague-Dawley rats (both male and female). Where indicated, we used male and female adult (7–14 weeks old) wild-type C57BL/6J mice (Beijing Vital River Laboratory and Charles River), *Dagla*-KO (C57BL/6J-*Dagla*^em1cyagen^; Cyagen), *Napepld*-KO (C57BL/6J-*Napepld*^em1cyagen^; Cyagen), and *Dagla*-flox (C57BL/6JCya-*Dagla*^em1flox/Cya^; Cyagen) mice. All animals were housed at 18–23 °C in 40–60% humidity under a standard 12 h/12 h light–dark cycle. Food and water were provided *ad libitum*.

### Drugs and reagents

For *in vitro* screening and sensor characterization, anandamide (AEA; MedChemExpress), 2-arachidonoylglycerol (2-AG; MedChemExpress), glutamate (Glu, Sigma-Aldrich), γ-aminobutyric acid (GABA, Tocris), glycine (Gly, Sigma-Aldrich), acetylcholine (ACh, Solarbio), 5-hydroxytryptamine (5-HT, Tocris), dopamine (DA, Sigma-Aldrich), norepinephrine (NE, Tocris), histamine (HA, Tocris), epinephrine (Epi, Sigma-Aldrich), adenosine (Ado, Tocris), ATP (Sigma-Aldrich), 1-oleoyl lysophosphatidic acid (LPA, Cayman), sphingosine-1-phosphate (S1P, Cayman), prostaglandin E2 (PGE2, Santa Cruz), prostaglandin D2 (PGD2, Cayman), prostaglandin F2α (PGF2α, Santa Cruz), ethanolamine (ETA, Aladdin), arachidonic acid (AA, Proteintech), nicotine (GLPBIO), cocaine (Qinghai Pharmaceutical Factory), Δ9-tetrahydrocannabinol (Δ9-THC), JWH-018, EG-018, ADB-BUTINACA, 4CN-CUMYL-BUTINACA, 5F-ADB, ADB-4en-PINACA and MDMB-4en-PINACA (Key Laboratory of Drug Monitoring and Control, Ministry of Public Security, China) were prepared as 10 mM stock solutions in ddH_2_O or DMSO. On the day of the experiments, these stock solutions were diluted to prepare a 10× working solution in Tyrode’s solution consisting of (in mM): 150 NaCl, 4 KCl, 2 MgCl_2_, 2 CaCl_2_, 10 HEPES, and 10 glucose (pH 7.4). A 10 μL aliquot of this working solution was then added to 90 μL Tyrode’s solution in each imaging well. For confocal and two-photon imaging using an imaging chamber, the stock solutions were diluted in Tyrode’s solution to a 3× working solution on the day of the experiment, and 500 μL of this working solution was added to 1000 μL Tyrode’s solution in the imaging chamber and mixed thoroughly. To maintain a constant volume, 500 μL of the mixed solution was then removed from the imaging chamber.

For acute brain slice experiments, AEA and 2-AG were prepared as 1000× stock solutions in aliquots and stored at −80 °C. On the day of the experiment, these stock solutions were diluted to their final concentrations in artificial cerebrospinal fluid (aCSF) containing (in mM): 130 NaCl, 26 NaHCO_3_, 10 glucose, 2.5 KCl, 2.5 CaCl_2_, 2 MgCl_2_, 1.25 NaH_2_PO_4_.

For *in vivo* recording experiments involving enzyme degradation inhibitors, JZL184 (MedChemExpress) and URB597 (MedChemExpress) were formulated in vehicle consisting of 10% DMSO, 40% PEG-300, 5% Tween-80, and 45% saline; where indicated, vehicle was used as a negative control.

For experiments involving addictive substances, nicotine, cocaine, ethanol, and MDMB-4en-PINACA were diluted in saline; Δ9-THC was prepared in a solution consisting of 5% DMSO, 5% Tween-80, and 90% saline.

### AAV expression

AAV2/9-hSyn-2-AG2.0 (7×10^13^ vg/mL) and AAV2/9-hSyn-AEA2.0 (2×10^13^ vg/mL) were packaged at WZ Biosciences. AAV2/9-hSyn-2-AG2.0 (1.2×10^13^ vg/mL), AAV2/9-hSyn-AEA2.0 (1.2×10^13^ vg/mL), and AAV2/9-hSyn-Cre (1.03×10^13^ vg/mL) were packaged at BrainVTA. Where indicated, the AAVs were either used to infect cultured neurons or injected *in vivo* into specific brain regions.

### Generation of AlphaFold3-predicted structural models and residue-ligand contact analysis

All structural models were predicted using a locally implemented version of AlphaFold3. For each protein complex, predictions were performed with the ‘num_diffusion_samples’ parameter set to 10 and the ‘modelSeeds’ parameter assigned a list of integers from 1 to 10, yielding 100 initial structures per CB1R-ligand complex (10 diffusion samples per seed). The recycle number was set as 0, 1, 2, and 3 for each of these structures, respectively. By combining these structural models with the default AlphaFold3 outputs, a total of 500 independent predicted structures were generated for each CB1R–ligand complex. Inputs to AlphaFold3 consisted of the CB1R sequence (PDB ID: 8ghv) and the SMILES strings of the respective ligands (2-AG/AEA).

For each set of 500 predicted structures, we first identified all amino acid residues located within 4 Å of any heavy atom of the bound ligands. The contact frequency between each such residue and the ligand was then calculated across all 500 structures (Fig. 1b). A contact was defined as a distance of ≤ 4 Å between any heavy atom of the ligand and any atom of the residue, a cutoff commonly used to characterize specific biomolecular interactions. From the set of 500 predicted structures, a residue was classified as a high-probability interaction site if it met both of the following frequency-based criteria: (1) it contacts count with a specific ligand heavy atom exceeded 200, and (2) it contacted at least two different heavy atoms of the ligand, each with a frequency greater than 100. These thresholds ensure that the identified interactions are stable and reproducible across the conformational ensemble.

### Molecular biology

Gibson assembly was used to generate all plasmids used in the study, and all sequences were verified using Sanger sequencing. For HEK293T cell selectivity screening and the sensor characterization experiments, coding sequences were inserted into the pDisplay vector (Invitrogen) fused with a downstream IRES-mCherry-CAAX cassette to serve as a cell membrane marker and to calibrate the sensor’s fluorescence. Site-directed mutagenesis was performed using primers synthesized by Tsingke Biological Technology, containing specific or randomized NNB codons. For viral packaging and neuronal expression, the sensors were cloned into a pAAV backbone under the control of the human synapsin (*SYN1*) promoter (pAAV-hSyn). For the luciferase complementation assay, WT-CB1R-SmBit and sensor-SmBit constructs were derived from the β2AR-SmBit template. For the Tango assay, genes encoding the WT receptor or the sensors were cloned into the pTango backbone.

### HEK293T cell culture and transfection

HEK293T cells (ATCC, CRL-3216) were maintained in high-glucose Dulbecco’s Modified Eagle’s Medium (DMEM; Gibco) supplemented with 10% (v/v) fetal bovine serum (FBS; CellMax) and 1% penicillin–streptomycin (Gibco). Cells were incubated at 37 °C in a humidified air containing 5% CO□. For sensor screening and characterization, cells were seeded into 96-well plates and allowed to reach approximately 70% confluence before transfection. Cells were transfected for 6–8 h using a mixture containing 0.3 μg plasmid DNA and 0.9 μg 40-kDa polyethylenimine (PEI) per well. For two-photon spectra measurements, cells were plated on 12-mm glass coverslips in 24-well plates and transfected with 1 μg DNA and 3 μg PEI per well for 6–8 h. Fluorescence imaging was performed 24–36 h after transfection.

### Generation of stable cell lines

To generate stable cell lines expressing the sensors, the sensor genes were cloned into the pPacific vector, which contains a 5′ terminal repeat, an IRES sequence, a puromycin resistance gene, and a 3′ terminal repeat.

HEK293T cells were seeded in 6-well plates and grown to 70%–80% confluence. The cells were then co-transfected with the pPacific plasmid containing the sensor gene and a PiggyBac transposase plasmid using PEI, with a plasmid:transposase:PEI mass ratio of 20:1:60; 36–48 h post-transfection, the culture medium was replaced with selection medium containing 1–3 μg/mL puromycin. The cells underwent multiple rounds of puromycin selection until uniformly expressing cell populations were established. These stably integrated cells were subsequently expanded in new culture dishes and validated for sensor expression and function prior to further experiments.

### Primary cortical neuron culture and viral transduction

Primary cortical neurons were isolated from P0 Sprague–Dawley rats (Beijing Vital River Laboratory). The cortices were dissected and dissociated with 0.25% trypsin–EDTA (Gibco), and the resulting cell suspension was plated onto 12-mm poly-D-lysine–coated glass coverslips (Sigma-Aldrich) in 24-well plates. The neurons were incubated at 37 °C in humidified air containing 5% CO_2_ in Neurobasal medium (Gibco) supplemented with 2% B-27 (Gibco), 1% GlutaMAX (Gibco), and 1% penicillin–streptomycin (Gibco); 50% of the culture medium was replaced every 3 days. At 3 days *in vitro* (DIV3), cytosine β-D-arabinofuranoside (Sigma) was added to a final concentration of 1 μM to limit glial cell proliferation. For sensor characterization in neurons, cortical neurons were infected at DIV3 with an AAV encoding the eCB sensors (full titer, 1 μL per well) and imaged at DIV17–24.

### Fluorescence imaging of cultured cells

Before imaging, the culture medium was replaced with Tyrode’s solution. The cells grown on coverslips were transferred to a custom-made imaging chamber and imaged using an inverted Ti-E A1 confocal microscope (Nikon) using NIS-Element 4.51.00 software (Nikon). The confocal microscope was equipped with a 10×/0.45 numerical aperture (NA) objective, a 20×/0.75 NA objective, a 40×/1.35-NA oil-immersion objective, a 488-nm laser, and a 561-nm laser. For experiments using neuronal electrical stimulation, the cells were loaded with the calcium dye Calbryte 590-AM (1 μM) for 30 mins prior to imaging.

The cells cultured in 96-well plates were imaged using either an Opera Phenix system (Perkin Elmer) equipped with a 20×/0.4-NA objective, a 40×/1.1-NA water-immersion objective, a 488-nm laser, and a 561-nm laser, or an Operetta CLS system equipped with a 20×/0.4-NA water-immersion objective, a 40×/1.1-NA water-immersion objective, a 488-nm LED, and a 561-nm LED controlled using Harmony 4.9 software.

The change in fluorescence (ΔF/F_0_) was calculated using the formula [(F–F_0_)/F_0_], in which F_0_ is baseline fluorescence defined as the average fluorescence measured 0–1 min before drug application.

### Spectra measurements

For measuring the one-photon spectra, cells stably expressing 2-AG2.0 or AEA2.0 were collected and transferred to 384-well plates in the absence or presence of 10 μM 2-AG or AEA, respectively. The excitation and emission spectra were then measured at 5 nm increments using a Safire2 multi-mode plate reader (Tecan). Non-transfected HEK293T cells were used for background subtraction.

For measuring the two-photon spectra, HEK293T cells at 50–60% confluence was transfected with pDisplay-CMV-2-AG2.0 or pDisplay-CMV-AEA2.0 and imaged 24 h post-transfection. The two-photon spectra of 2-AG2.0 and AEA2.0 were measured at 10 nm increments ranging from 700 to 1050 nm using an Ultima Investigator two-photon microscope (Bruker) equipped with a 20×/1.00-NA water-immersion objective (Olympus), an InSight X3 tunable laser (Spectra-Physics), and Prairie View 5.5 software (Bruker). Non-transfected cells were used for background subtraction. Laser power was normalized according to the output power of the tunable laser with different wavelengths.

### Luciferase complementation assay

Upon reaching 50–60% confluence, HEK293T cells were co-transfected with the LgBit-mG construct and either the WT receptor or relevant sensor; 24–36 h post-transfection, the cells were mechanically harvested using a scraper, resuspended in phosphate-buffered saline (PBS), and transferred to 96-well plates. The cells were then treated with 5 μM furimazine (NanoLuc Luciferase Assay; Promega) and varying concentrations of 2-AG or AEA (ranging from 0.01 nM to 100 μM). Following a 10-min incubation period in the dark at room temperature, luminescence was quantified using a VICTOR X5 multi-label plate reader (PerkinElmer).

### Tango assay

Plasmids encoding 2-AG2.0, AEA2.0, or WT CB1R were transfected in a specialized reporter cell line that stably expresses a β-arrestin 2–TEV fusion protein alongside a tTA-dependent luciferase reporter gene (HTLA cell line, a gift from Bryan Roth lab). Twenty-four hours post-transfection, the cells were collected using trypsin digestion and re-seeded in 96-well plates. Agonist stimulation was initiated by applying AEA or 2-AG at concentrations ranging from 0.01 nM to 10 μM. Following a 12-h incubation period to allow for luciferase expression, the assay was developed by adding Bright-Glo (Fluc Luciferase Assay System; Promega) to a final concentration of 5 μM, and luminescence was quantified using a VICTOR X5 multi-label plate reader (PerkinElmer).

### Fluorescence imaging of acute striatal slices

C57BL/6J mice (7–10 weeks old) were anesthetized with isoflurane and placed in a small animal stereotaxic frame (David Kopf Instruments). After exposing the skull under aseptic techniques, small burr holes were drilled, and AAV solutions were injected bilaterally (1 µL/site at a rate of 200 nL/min) through a 32-gauge syringe (Hamilton) using a microsyringe pump (World Precision Instruments). AAV9-hSyn-2AG2.0 (1.2×10^13^ vg/mL), and AAV9-hSyn-AEA2.0 (1.2×10^13^ vg/mL) were injected into the dorsolateral striatum (DLS) using the following coordinates: AP, +0.8 mm relative to Bregma; ML, ±1.8 mm relative to Bregma; and DV, −2.4 mm from the dura.

Acute striatal slices were prepared 3–4 weeks after AAV injection using standard techniques. In brief, the mice were sacrificed by cervical dislocation, and the brains were dissected out and submerged in ice-cold cutting solution consisting of (in mM): 194 sucrose, 30 NaCl, 26 NaHCO_3_, 10 glucose, 4.5 KCl, 1.2 NaH_2_PO_4_, 1 MgCl_2_. Coronal slices (300-µm thick) containing the striatum were cut using a vibratome (VT1200S, Leica Microsystems) and transferred to a holding chamber containing aCSF. The slices were incubated at 34 °C for 15 min before being stored at room temperature until recordings were performed. All solutions were saturated with 95% O_2_/5% CO_2_.

Individual slices were hemisected and transferred to a recording chamber that was superfused at ∼2 mL/min with aCSF at 31–33 °C. A microscope (SliceScope, Scientifica) equipped with a 470-nm LED (2.8 mW/mm^2^; pE-300, CoolLED), CMOS camera (Prime BSI Express, Teledyne Photometrics), 525/50-nm emission filter (Cairn Research), and 10×/0.3-NA water-immersion objective (Olympus) was used for wide-field fluorescence imaging of 2-AG2.0 and AEA2.0. Image acquisition was controlled using Micro-Manager1.4. Electrical stimulation, LED light application, and image acquisition were synchronized using TTL-driven stimuli via Multi Channel Stimulus II (Multi Channel Systems). Image files were analyzed using Fiji1.5 and MATLAB R2017b. For experiments measuring changes in basal, non-stimulated extracellular eCB levels, images (exposure duration: 100 ms) were acquired every 10 s for 50 min. Mean fluorescence intensity (F) was extracted from a region of interest (ROI) 100×100 µm. Data are expressed as the change in fluorescence (ΔF/F_0_) and were calculated using the formula [(F–F_0_)/F_0_], where F_0_ is the average F measured during the initial 10 min. For experiments measuring extracellular eCB levels in response to trains of electrical stimuli, images were acquired at 10 Hz (exposure duration: 100 ms) for 20 s. Electrical pulses (pulse duration: 200 µs; pulse amplitude: 0.6 mA) were delivered via a local bipolar concentric Pt/Ir electrode (inner diameter: 25 µm; outer diameter: 125 µm; FHC). F was extracted from a 100×100 µm ROI approximately 100 µm from the stimulating electrode, and ΔF/F_0_ was calculated using the formula [(F–F_0_)/F_0_], where F_0_ was determined by fitting a two-term exponential decay function [f(*x*) = a*e*^b*x*^ + c*e*^d*x*^] to pre-peak F.

### *In vivo* fiber photometry recordings in mice

#### Foot shock

Adult (8–10 weeks old) WT C57BL/6J, *Dagla*-KO, and *Napepld*-KO mice were anesthetized, and 300 nL of AAV9-hSyn-2-AG2.0 (7×10^13^ vg/mL) or AAV9-hSyn-AEA2.0 (2×10^13^ vg/mL) was injected bilaterally into either the basolateral amygdala (BLA) using the following coordinates: AP, −1.4 mm relative to Bregma; ML, ±3.0 mm relative to Bregma; and DV, −4.2 mm from the dura or the nucleus accumbens (NAc) core using the following coordinates: AP, +1.4 mm relative to Bregma; ML, ±1.2 mm relative to Bregma; and DV, −4.3 mm from the dura. An optical fiber (200-μm core diameter, 0.39-NA; RWD) was implanted 0.1 mm dorsal to the virus injection site in the BLA or NAc core and secured to the skull using dental cement.

For the fear-related foot-shock experiments, the mice were tested in a conditioned fear system (Med Associates). The mice were first habituated to the chamber and fiber photometry tethering for 3 consecutive days before the experiment to minimize stress related to the environment and cable attachment. Fiber photometry recordings were performed using an R821 system (RWD); 410 nm and 470 nm LEDs were used to excite the sensors. During the recording session, mice were placed in the chamber for a 3-min baseline period, followed by five 2-s foot shocks (0.5 mA), with a 180-s interval between each shock.

Fiber photometry data from the foot-shock experiments were analyzed using OFRS software (RWD). Photobleaching was corrected using partial least squares fitting, and motion- and/or hemodynamics-related artifacts were minimized using the 470 nm excitation signal corrected with the 410 nm excitation channel. The baseline was defined as mean fluorescence measured during the 0–5 s period immediately preceding the foot shocks. To compare fluorescence changes across animals, the ΔF/F_0_ traces were normalized by the standard deviation of the baseline, yielding *z*-score signals. The area under the curve (AUC) and the rise and decay times (10–90%) were quantified using OriginPro 2024.

#### Addictive substances administration

Adult (8–10 weeks old) C57BL/6J and *Napepld*-KO mice were anesthetized, and 300 nL of AAV9-hSyn-2-AG2.0 (7×10^13^ vg/mL) or AAV9-hSyn-AEA2.0 (2×10^13^ vg/mL) was injected bilaterally into the nucleus accumbens (NAc) shell using the following coordinates: AP, +1.4 mm relative to Bregma; ML, ±0.9 mm relative to Bregma; and DV, −4.5 mm from the dura. For the adult *Dagla*-flox mice, AAV9-hSyn-2-AG2.0 was mixed at a 1:1 ratio with AAV9-hSyn-Cre (matched titer), and a total volume of 500 nL was injected bilaterally into the NAc shell using the coordinates above. An optical fiber (200-μm core diameter, 0.39-NA; RWD) was implanted 0.1 mm dorsal to the virus injection site in the NAc shell and secured to the skull using dental cement.

The animals were habituated to the open-field recording chamber and fiber photometry tethering for 3 consecutive days prior to the experiment. Fiber photometry recordings were acquired using an R821 system (RWD); a 470 nm LED was used for sensor excitation. For the nicotine (0.5 mg/kg), cocaine (20 mg/kg), ethanol (2 g/kg, 40% solution), and MDMB-4en-PINACA (0.1 mg/kg) experiments, the mice were placed in the chamber for a 30-min baseline period followed by 2-h of continuous recording after drug injection. For the Δ9-THC experiments, a 1-h baseline period was used, followed by at least 10 h of recording after drug injection.

Fiber photometry data were processed and analyzed using OFRS software (RWD), Spike2 (CED), and custom Python scripts. The baseline was defined as mean fluorescence measured during the period preceding drug injection. Data acquired during the 5-min injection window was excluded from analysis due to large motion artifacts. ΔF/F_0_ traces were then normalized by the standard deviation of the baseline to yield *z*-score signals, enabling comparison of fluorescence changes across animals. The AUC and the rise and decay times (10–90%) were quantified using OriginPro 2024 and custom-written Python scripts. The locomotion videos of the animal were analyzed by EzTrack (https://github.com/denisecailab/eztrack) and custom-written Python scripts.

### Locomotor analysis

Locomotor speed was extracted from the open-field videos using EzTrack and custom Python scripts. For each mouse, relative speed was calculated by dividing the speed at each time point by the mean speed measured during the pre-injection baseline window (−65 to −5 min). The relative-speed traces were smoothed using a Savitzky–Golay filter and constrained to non-negative values. Normalized moving distance was calculated as the discrete time integral of the relative-speed trace:

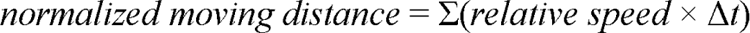

Because relative speed is dimensionless, the integrated values are reported in arbitrary units (A.U.) rather than meters. The Δ9-THC induced locomotion change was calculated from 5 to 125 min after injection.

### Immunohistochemistry

Following the experiments, the mice were deeply anesthetized and transcardially perfused with PBS, followed by 4% paraformaldehyde (PFA) in PBS. The brains were dissected and post-fixed overnight in 4% PFA at 4 °C, then cryoprotected in 30% sucrose in PBS until saturated. Coronal sections were cut at 40-μm thickness using a cryostat microtome (CM1950; Leica). The sections were incubated in blocking solution containing 5% (v/v) normal goat serum, 0.1% Triton X-100, and 2 mM MgCl_2_ in PBS for 1 h at room temperature, followed by overnight incubation at 4 °C in AGT solution (0.5% normal goat serum, 0.1% Triton X-100, and 2 mM MgCl_2_ in PBS) containing chicken anti-GFP primary antibody (10 mg/mL, 1:500; Cat# ab13970, Abcam). The following day, the sections were rinsed three times in AGT solution and incubated for 2 h at room temperature with Alexa Fluor 488-conjugated goat anti-chicken secondary antibody (2 mg/mL, 1:500; Cat# ab150169, Abcam); the nuclei were counterstained with DAPI (5 mg/mL, 1:1,000; Cat# HY-D0814, MedChemExpress). After three additional washes in AGT solution, the sections were mounted on glass slides and imaged using a VS120-S6-W Virtual Slide Microscope (Olympus) equipped with a 10×/0.4-NA objective.

## Quantification and statistical analysis

All data were analyzed and plotted using Microsoft Excel, OriginPro 2024, Spike2, custom-written MATLAB or Python scripts, where appropriate. Unless stated otherwise, summary data are presented as the mean ± the standard error of the mean (SEM). Groups were compared using a two-tailed Student’s *t*-test or one-way analysis of variance (ANOVA), where appropriate. For multiple comparisons following ANOVA, Tukey’s or Dunnett’s post-hoc tests were applied. Significance levels are denoted in the figures as follows: \**p* < 0.05, \*\**p* < 0.01, \*\*\**p* < 0.001, and n.s., not significant (*p* > 0.05).

## Acknowledgments

This research was supported by grants from the National Key R&D Program of China (2024YFF1206400 to H.W., 2023YFE0207100 to Y.L., 2022YFC3300905 to H.D.); the National Science and Technology Innovation 2030-Major Project of China (Grant No. STI2030-Major Projects 2022ZD0208300 to Z.W. and R.C.); the National Natural Science Foundation of China (32525003 and 31925017 to Y.L, W2542016 to W.T.), Beijing Municipal Science & Technology Commission (Z220009 to Y.L.); grants from the Feng Foundation of Biomedical Research, the New Cornerstone Science Foundation through the New Cornerstone Investigator Program; grants from the Peking-Tsinghua Center for Life Sciences and the State Key Laboratory of Membrane Biology at Peking University School of Life Sciences (to Y.L), and grants from Aligning Science Across Parkinson’s (ASAP) through the Michael J. Fox Foundation for Parkinson’s Research (ASAP-020370 and ASAP-025192 to S.C.).

We thank the optical imaging platform and small animal imaging platform of the National Center for Protein Sciences at Peking University in Beijing, China, for their support and assistance with the Operetta CLS high-content imaging system and the behavior facility; the Laboratory Animal Center of Peking University for advice and technical support; X. Lei at PKU-CLS for assistance with the Opera Phenix high-content screening system. We thank members and alumni of the Yulong Li group for support and discussions. Some diagrams were created using BioRender.com.

## Author Contributions

Y.L. conceived the project. R.C., S.Cai., W.T., and A.D. performed the experiments for sensor development, optimization and *in vitro* characterization. R.C. and Y.Y. performed the *in vivo* fiber photometry recordings and related experiments. A.S.S. and K.L.T. performed the acute brain slice imaging experiments under the supervision of S.J.C. R.C. performed the *in silico* prediction experiment, and S.Chen., L.W., and C.S. provided assistance on the *in silico* prediction strategies. P. X., Y.Q. and H.D. provided assistance on the Δ9-THC and synthetic cannabinoids experiments; Z.W. and H.W. contributed to manuscript revision; R.C. and Y.L. wrote the manuscript with input from all other authors.

---

**Supplementary Fig. 1:**
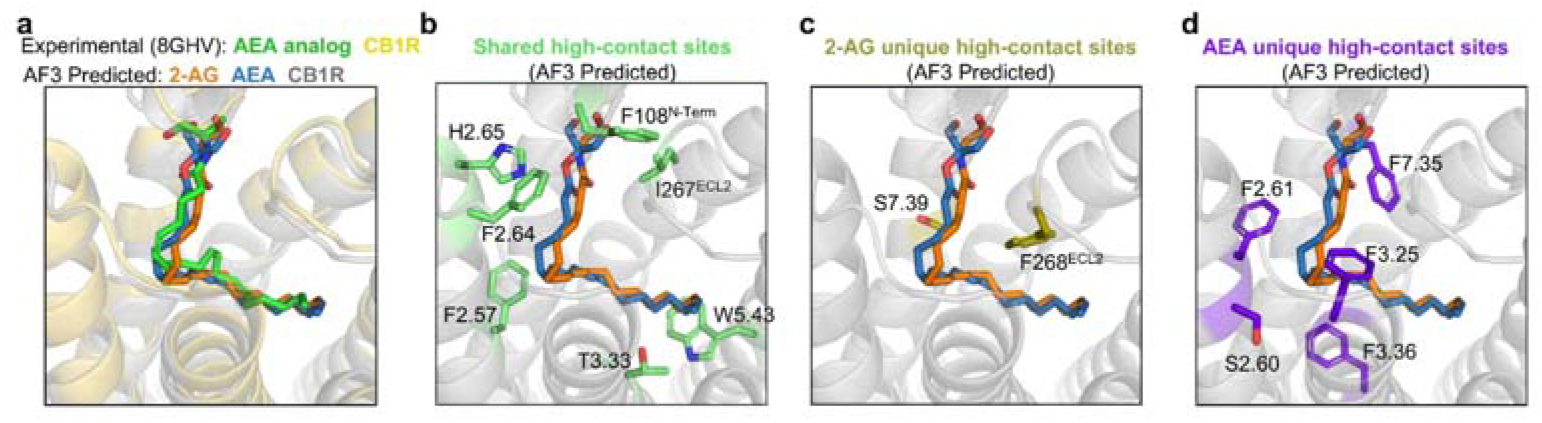
High-contact-frequency residues of CB1R involved in 2-AG and AEA binding, as identified from AlphaFold3-predicted structures. a. Structural alignment of the experimentally determined CB1R–AMG315 (AEA analog) complex (PDB: 8GHV) with the AlphaFold3-predicted CB1R–2-AG and –AEA binding structures. b–d. AlphaFold3-predicted structures of wild-type CB1R in complex with 2-AG or AEA. Residues with high contact frequency that are shared between both ligands (b), unique to 2-AG (c), or unique to AEA (d) are highlighted.

**Supplementary Fig. 2:**
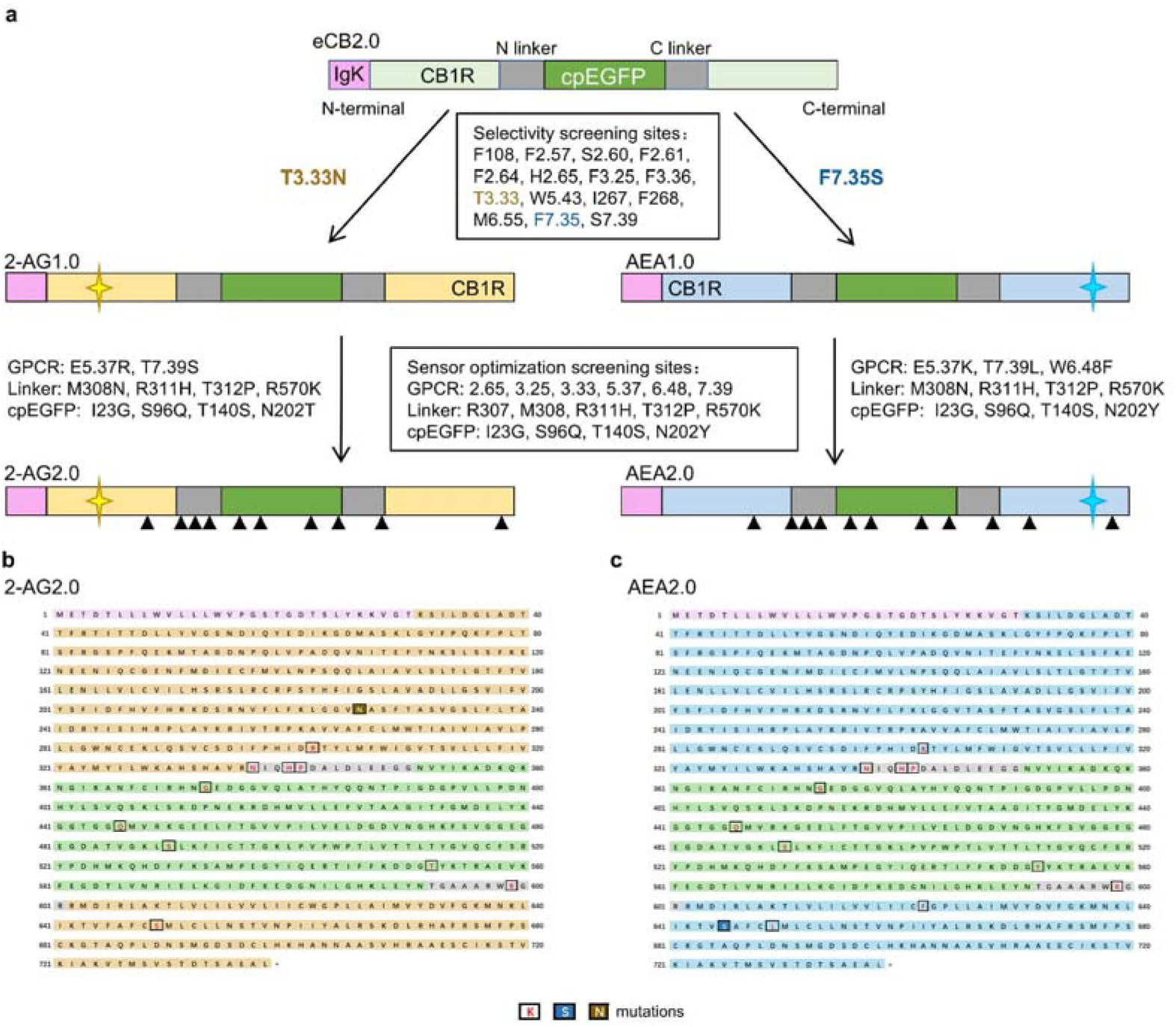
Strategy for developing the GRAB_2-AG_ and GRAB_AEA_ sensors. a. Diagram illustrating the process for developing the 2-AG2.0 (left) and AEA2.0 (right) sensors. The triangles indicate the approximate locations of the indicated mutations. b, c. The full amino acid sequences of the 2-AG2.0 (b) and AEA2.0 (c) sensors. The IgK leader sequence (pink), receptor backbone (yellow for 2-AG2.0, blue for AEA2.0), linkers (gray), and cpEGFP (green) are highlighted. The key mutations introduced to generate the selective sensors, namely T3.33N in 2-AG1.0 (b) and F7.35S in AEA1.0 (c), are labeled in dark blue and brown, respectively. Other residues to generate the final sensors that were mutated during the optimization process are also indicated in red.

**Supplementary Fig. 3:**
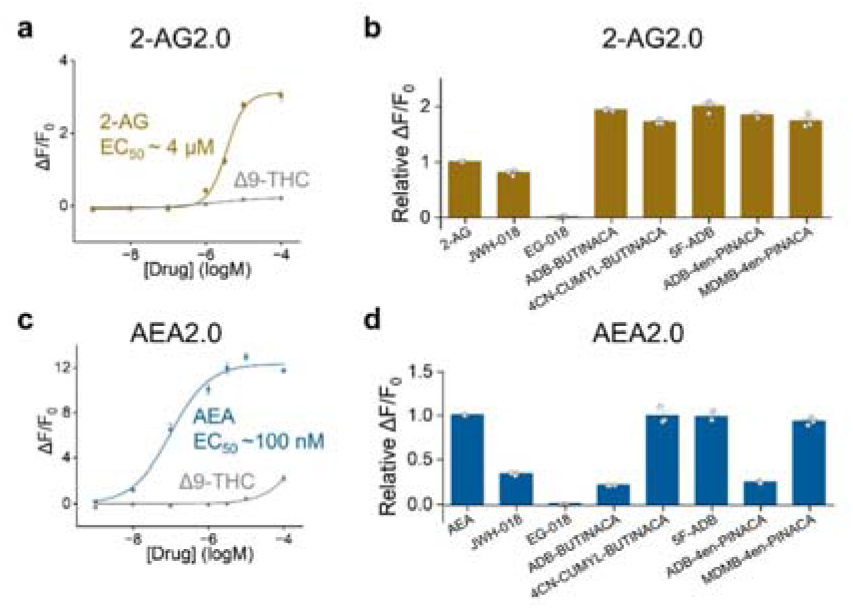
Response of 2-AG2.0 and AEA2.0 to synthetic cannabinoids and phytocannabinoids. a, c. Dose–response curves for 2-AG2.0 (a) and AEA2.0 (c) in response to Δ9-THC, and their respective ligands (AEA or 2-AG). The calculated EC_50_ values are indicated in the plots. b, d. Summary of the fluorescence response measured in HEK293T cells expressing 2-AG2.0 (b) or AEA2.0 (d) following application of the indicated endocannabinoids (2-AG or AEA) and various synthetic cannabinoids. All responses are relative to the maximum response elicited by 2-AG (for 2-AG2.0) or AEA (for AEA2.0).

**Supplementary Fig. 4:**
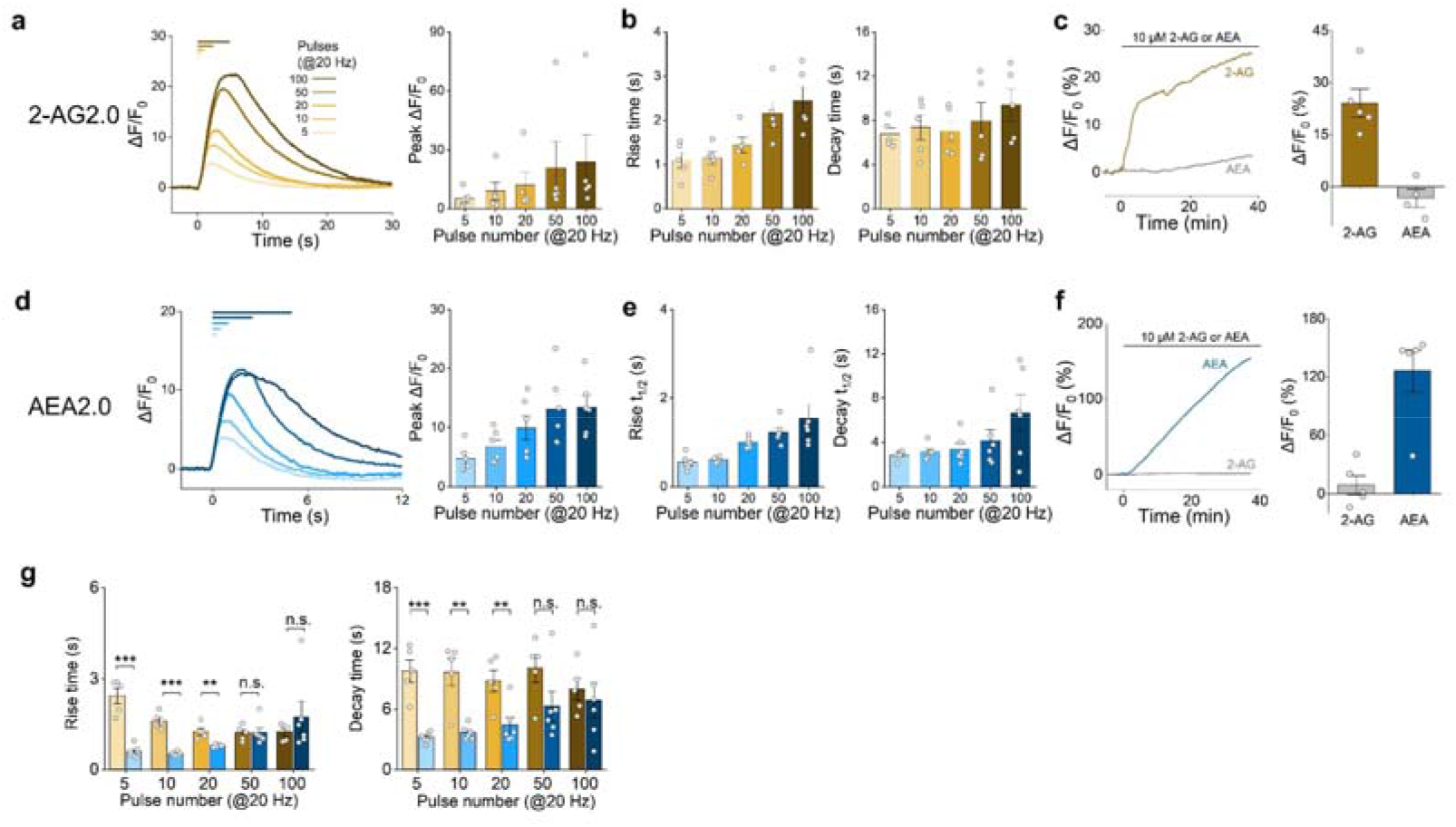
Characterization of the 2-AG2.0 and AEA2.0 sensors in acute brain slices. a, d. Representative fluorescence traces (left) and summary of the peak signals (right) measured in brain slices expressing 2-AG2.0 (a) or AEA2.0 (d) before and after the indicated electrical stimuli; n = 5–6 slices per group. b, e. Summary of the rise (left) and decay (right) times measured for the 2-AG2.0 (b) and AEA2.0 (e) fluorescence signals in response to the indicated number of electrical pulses applied at 20 Hz; n = 5–6 slices per group. c, f. Representative fluorescence traces (left) and summary of the maximum signals (right) measured in brain slices expressing 2-AG2.0 (c) or AEA2.0 (f) before and during the application of 2-AG or AEA; n = 4–5 slices per group. g. Rise and decay times measured for the 2-AG2.0 (yellow) and AEA2.0 (blue) fluorescence signals in response to the indicated number of electrical pulses applied at 20 Hz. These data are replotted from (b) and (e) for comparison purposes only.

**Supplementary Fig. 5:**
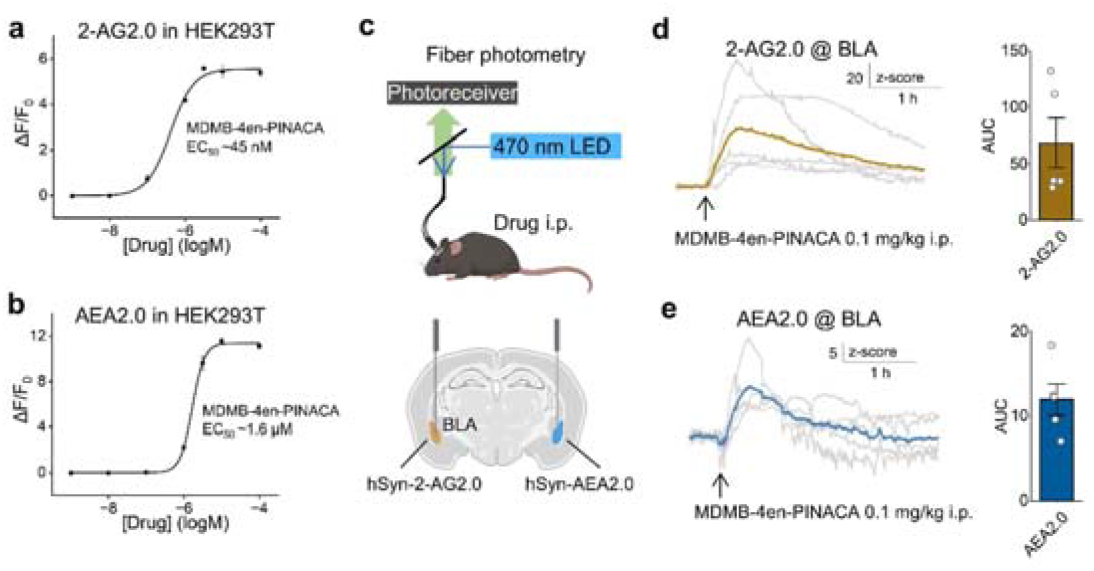
Control experiments confirming expression of 2-AG2.0 and AEA2.0 in the BLA of WT mice. a, b. Dose–response curves for 2-AG2.0 (a) and AEA2.0 (b) in response to MDMB-4en-PINACA. The calculated EC_50_ values are indicated in the plots. c. Schematic diagram depicting the experimental strategy for *in vivo* fiber photometry recording and the injection of AAVs expressing 2-AG2.0 and AEA2.0 in the BLA of opposing hemispheres, with fiber implantation. d, e. Left: average fluorescence traces (*z*-score) of 2-AG2.0 (d) and AEA2.0 (e) in the BLA before and after an i.p. injection (indicated by the arrow) of MDMB-4en-PINACA (0.1 mg/kg). Right: summary of the area under the curve (AUC) for the 2-AG2.0 (d) and AEA2.0 (e) responses.

**Supplementary Fig. 6:**
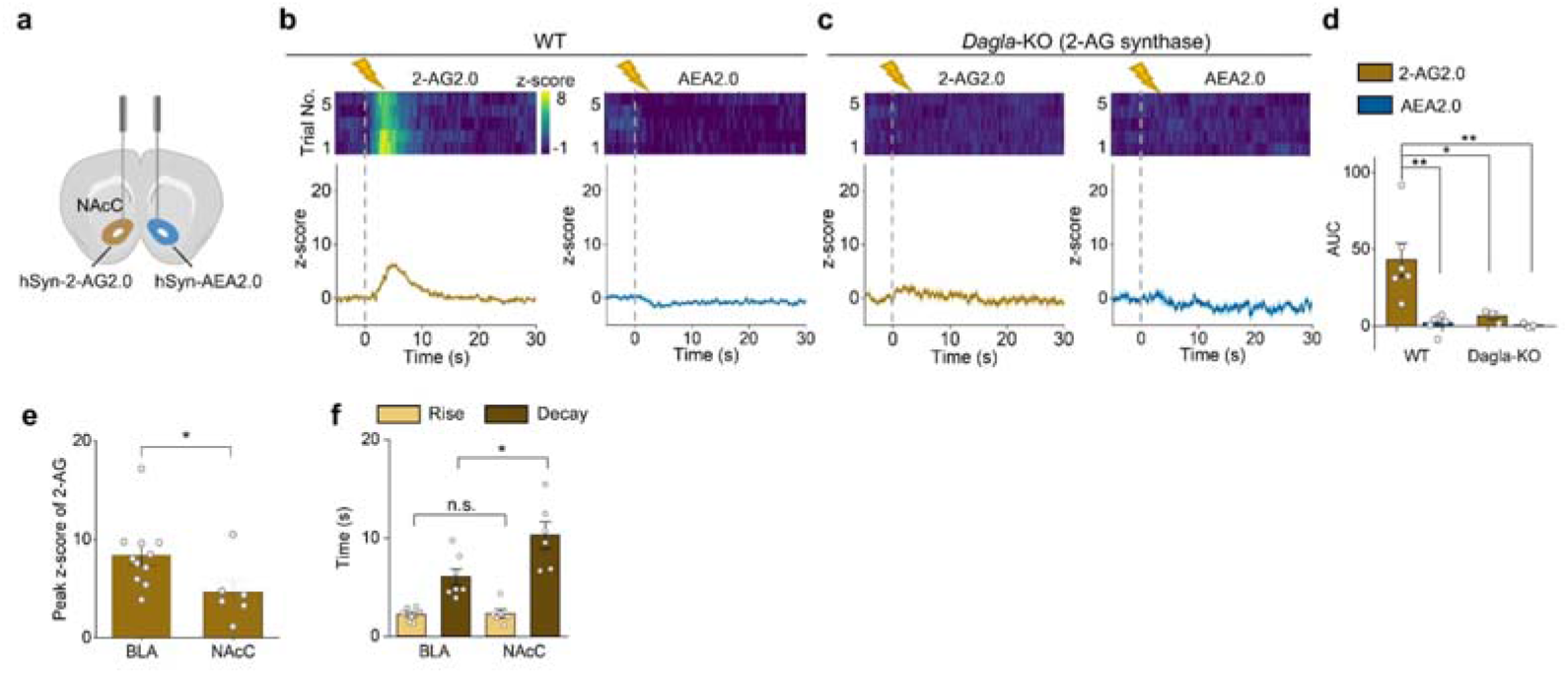
2-AG and AEA dynamics measured in the NAc core in response to foot shock. a. Schematic diagram showing the injection of AAVs expressing 2-AG2.0 and AEA2.0 in the NAc core (NAcC) of opposing hemispheres, with fiber implantation. b, c. Representative pseudocolor heatmaps showing sensor responses over 5 trials (top) and corresponding average fluorescence traces (bottom) measured in the NAc core of a WT mouse (b) and a *Dagla*-KO mouse (c) expressing 2-AG2.0 (yellow traces) and AEA2.0 (blue traces) in opposing hemispheres. Where indicated, a 2-s foot shock was applied. d. Summary of the area under the curve (AUC) of the 2-AG2.0 and AEA2.0 fluorescence signals measured in the NAc core of WT and *Dagla*-KO mice following foot shock; n = 3–6 mice per group. e. Summary of the peak *z*-score for the foot shock-evoked 2-AG2.0 signals measured in the BLA (replotted from Fig. 4f) and NAc core; n = 6–11 mice per group. f. Summary of the rise and decay times (10–90%) for the foot shock-evoked 2-AG2.0 signals measured in the BLA (replotted from Fig. 4f) and NAc core; n = 6–11 mice per group.

**Supplementary Fig. 7:**
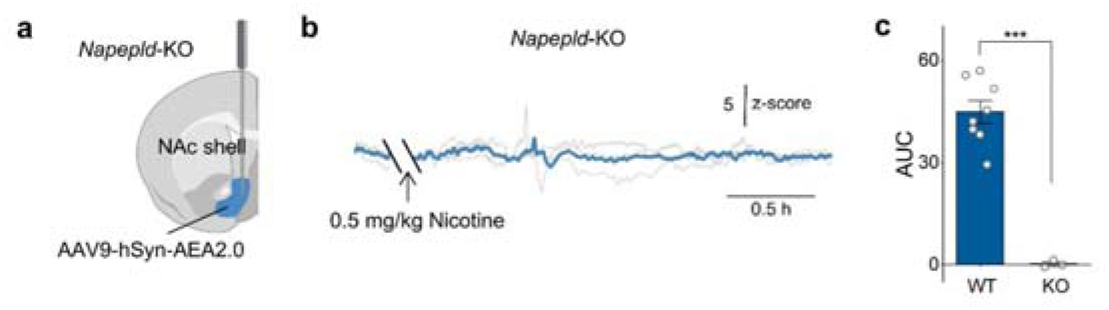
Validation of the nicotine-evoked AEA signal in the NAc shell using *Napepld*-KO mice. a, Schematic depicting AEA2.0 expression and optical-fiber implantation in the NAc shell of *Napepld*-KO mice. b. Average AEA2.0 fluorescence trace (*z*-score) recorded in the NAc shell of Napepld-KO mice before and after i.p. injection of nicotine (0.5 mg/kg). The arrow indicates the time of injection; gray lines represent individual mice and the blue line represents the group mean. c, AUC of the AEA2.0 signal from 5 to 35 min after nicotine injection in WT (replotted from Fig. 5d2) and Napepld-KO mice. n = 3–9 mice per group.

**Supplementary Fig. 8.**
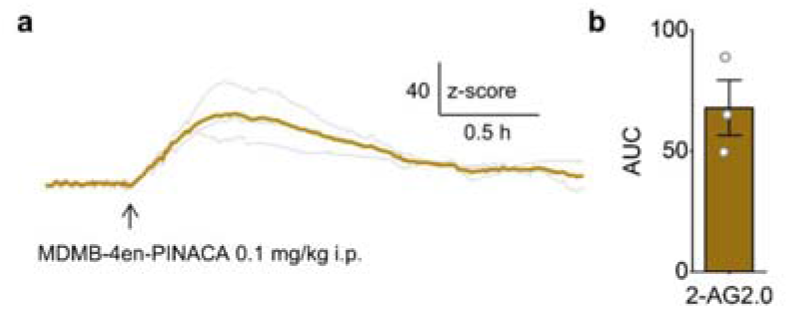
Control experiment confirming 2-AG2.0 expression and responsiveness in the NAc shell of *Dagla*-cKO mice. a, Average 2-AG2.0 fluorescence trace (*z*-score) recorded in the NAc shell of *Dagla*-cKO mice before and after i.p. injection of MDMB-4en-PINACA (0.1 mg/kg). The arrow indicates the time of injection; gray lines represent individual mice and the yellow line represents the group mean. b. AUC of the 2-AG2.0 signal from 5 to 35 min after injection; n = 3 mice.

